# Paresthesia during spinal cord stimulation depends on synchrony of dorsal column axon activation

**DOI:** 10.1101/2023.01.10.523167

**Authors:** Boriss Sagalajev, Tianhe Zhang, Nooshin Abdollahi, Noosha Yousefpour, Laura Medlock, Dhekra Al-Basha, Alfredo Ribeiro-da-Silva, Rosana Esteller, Stéphanie Ratté, Steven A. Prescott

## Abstract

Spinal cord stimulation (SCS) reduces chronic pain. Conventional (40-60 Hz) SCS engages spinal inhibitory mechanisms by activating low-threshold mechanoreceptive afferents with axons in the dorsal columns (DCs). But activating DC axons typically causes a buzzing sensation (paresthesia) that can be uncomfortable. Kilohertz-frequency (1-10 kHz) SCS produces analgesia without paresthesia and is thought, therefore, not to activate DC axons, leaving its mechanism unclear. Here we show in rats that kilohertz-frequency SCS activates DC axons but causes them to spike less synchronously than conventional SCS. Spikes desynchronize because axons entrain irregularly when stimulated at intervals shorter than their refractory period, a phenomenon we call overdrive desynchronization. Effects of overdrive desynchronization on evoked compound action potentials were verified in simulations, rats, pigs, and a chronic pain patient. Whereas synchronous spiking in DC axons is necessary for paresthesia, asynchronous spiking is sufficient to produce analgesia. Asynchronous activation of DC axons thus produces paresthesia-free analgesia.

## INTRODUCTION

Inhibitory interneurons in the spinal dorsal horn regulate transmission of pain signals to the brain^1^. Low-threshold mechanoreceptive afferents activate spinal inhibitory interneurons via axon collaterals that enter the spinal grey matter while also conveying touch signals via their main axon in the ipsilateral DC. In contrast, high-threshold afferents (nociceptors) terminate in the spinal dorsal horn, leaving pain signals to be relayed by second-order neurons that ascend contralaterally in anterolateral tracts. Anatomical separation of the pathways mediating touch and pain provides an ideal opportunity to selectively stimulate the former in order to inhibit the latter^2^. Just two years after gate control theory was proposed, SCS targeting the DCs was shown to reduce pain^3^. Though gate control mechanisms do not fully explain SCS effects, spikes conducted antidromically in DC axons modulate pain signals in the dorsal horn^4^ while spikes conducted orthodromically evoke a buzzing sensation known as paresthesia. Paresthesia is useful for targeting SCS^5^ and, until recently, was considered necessary for SCS-mediated pain relief^4, 6^. But paresthesia can be uncomfortable at high SCS intensities^7^ (especially when perception threshold is decreased by postural changes^8, 9^), thus limiting the strength of SCS and its therapeutic efficacy^10^. Discovery of paresthesia-free (subperception) forms of SCS is therefore clinically significant^11-14^.

One form of subperception SCS involves stimulating tonically at 1-10 kHz, instead of the 40-60 Hz used for conventional SCS (c-SCS). Absence of paresthesia during kilohertz-frequency SCS (kf-SCS) suggests – and early experiments^15^ and simulations^16^ seemingly confirmed – that DC axons are not activated, prompting a search for novel mechanisms. Subsequent experiments showed that DC axons can be activated by kf-SCS but did not explain the lack of paresthesia^17^. Direct effects of kf-SCS on dorsal horn cells (i.e. not via DC axons) has garnered attention^18^ but direct inhibition was shown to be artefactual^19^ and direct activation is incompatible with dorsal horn neurons having a higher activation threshold than DC axons^20^. There is good evidence that kf-SCS ultimately activates inhibitory neurons^21, 22^ and induces synaptic depression^23^ in the spinal dorsal horn, but how it does so – if not by activating DC axons – remains unclear.

Starting with experiments in rats, we show that kf-SCS, like c-SCS, activates DC axons. The critical difference is that c-SCS evokes action potentials (spikes) that are synchronized across axons whereas kf-SCS evokes asynchronous spikes. Spikes desynchronize when axons are stimulated at intervals shorter than their refractory period, a mechanism we call overdrive desynchronization. We also show that evoked compound action potentials (ECAPs), whose amplitude is correlated with paresthesia intensity, are attenuated by overdrive desynchronization. Whereas synchronous spikes are necessary for paresthesia, asynchronous spikes are sufficient to produce analgesia. Controlling spike synchrony is critical for maximizing pain relief by minimizing uncomfortable somatosensory percepts during SCS or, conversely, for deliberately producing somatosensory percepts in other applications, like neural prostheses^24^.

## RESULTS

### Modulation of spinal reflexes and cortical activity

SCS was applied to healthy rats under urethane anesthesia via a preclinical lead positioned midline at the T13 spinal level (**Fig. 1A**). Constant-current, symmetrically biphasic 40 µs-long pulses were delivered using a narrow (1 mm) bipole. Motor threshold (MT) was determined as the lowest pulse amplitude at which 2 Hz SCS caused muscle twitches (**Supplementary Fig. 1**). Sustained contraction occurred during c-SCS at intensities ≥MT, like in human studies^25^. Consistent with past reports^26-28^, transient contraction occurred at the onset of high-intensity kf-SCS but could be prevented by ramping up to the final intensity; this onset response occurred at intensities as low as 60% MT. Unless otherwise indicated, c-SCS and kf-SCS were tested at ∼50% MT without ramping.

**Figure 1.**
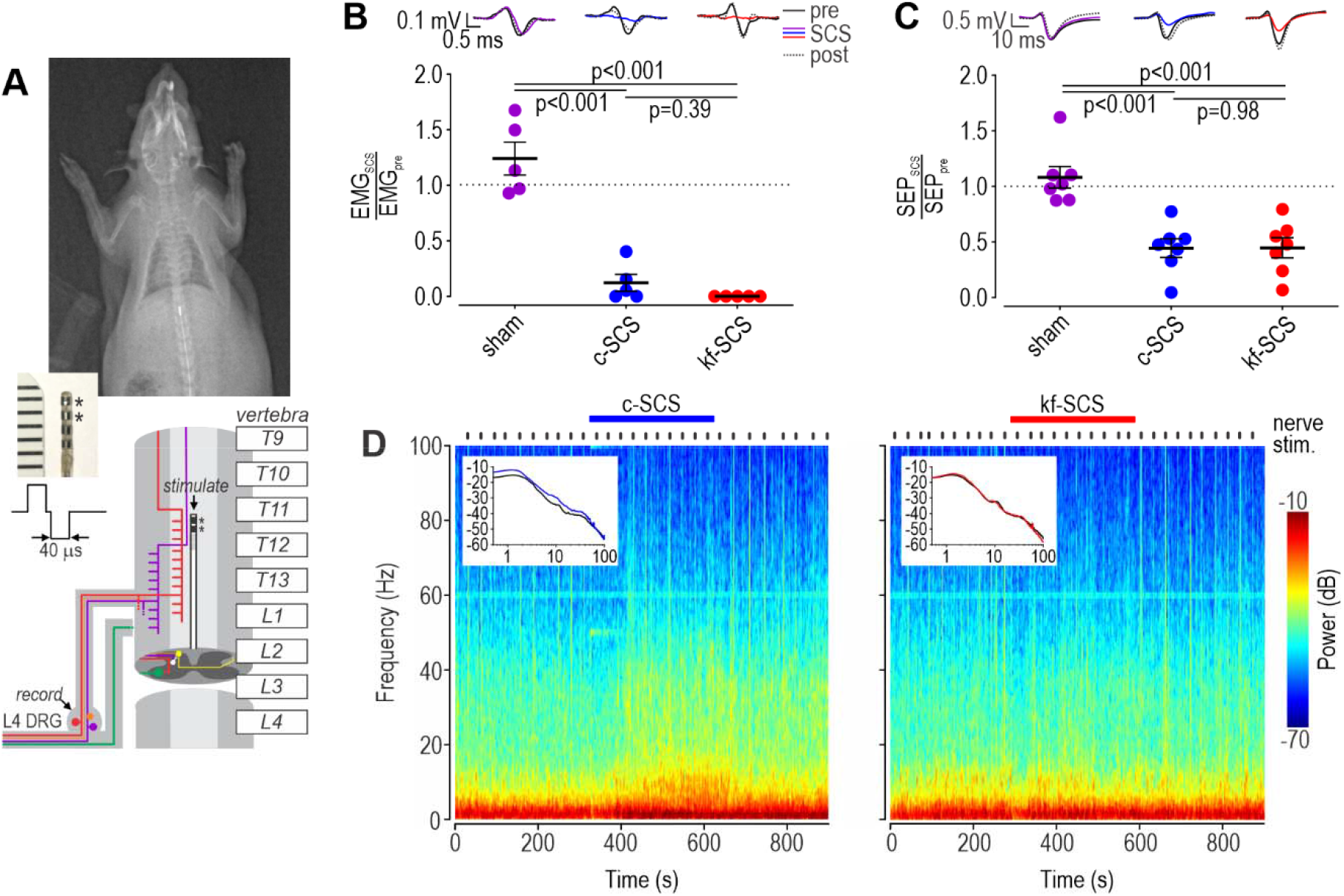
Modulation of spinal reflexes and cortical activity by SCS in rats. **(A)** Lead placement. Cartoon shows routes of different afferent types – mechanoreceptors (purple), proprioceptors (red) and nociceptors (orange) – plus motor neurons (green) and dorsal horn neurons (white, yellow). A symmetrically biphasic waveform was applied using contacts marked with *s. **(B)** H reflex was significantly attenuated by SCS (*F*_2,12_=50.84, p<0.001, one-way ANOVA); post-hoc Student-Neuman-Keuls (SNK) tests are reported on the graph. Traces show sample electromyogram (EMG) traces before (solid black), during (color) and after (dotted black) sham SCS (purple), c-SCS (blue) or kf-SCS (red). Data are plotted as the ratio of response amplitude before vs during SCS, and summarized as mean±SEM. **(C)** Somatosensory evoked potential (SEP) was significantly attenuated by SCS (*F*_2,12_=16.39, p<0.001, one-way ANOVA). Data are presented and analyzed like in B. **(D)** Ongoing cortical activity was altered by c-SCS but not by kf-SCS according to spectrograms. Insets show power spectra for epoch before (black) and during (color) SCS. Sciatic nerve stimuli are indicated along top and resulting SEPs are evident on the spectrogram as thin vertical strips reflecting transient cortical activation.

Since previous studies already established that kf-SCS, like c-SCS, has analgesic effects in rats^15, 28^, we focused on other clinically used tests^29^ in which c-SCS and kf-SCS have not been compared. To assess modulation at the spinal level, we triggered the H reflex via electrical stimulation of the medial plantar nerve and compared the electromyogram (EMG) H wave before, during, and after SCS. The H wave was significantly attenuated by c-SCS (*Q*=11.67, *p*<0.001, Student-Newman-Keuls [SNK] test) and by kf-SCS (*Q*=12.93, *p*<0.001) to a similar degree (*Q*=1.26, *p*=0.39) (**Fig. 1B**). To assess modulation at the cortical level, we electrically stimulated the sciatic nerve and measured the somatosensory evoked potential (SEP) from primary somatosensory (S1) cortex. Like the H reflex, the SEP was significantly attenuated by c-SCS (*Q*=7.03, *p*<0.001, SNK test) and by kf-SCS (*Q*=6.99, *p*< 0.001) to a similar degree (*Q*=0.034; *p*=0.98) (**Fig. 1C**). Separate from the transient activation caused by intermittent nerve stimulation, ongoing cortical activity was increased by c-SCS but not by kf-SCS (**Fig. 1D**), which may correlate with paresthesia and reflect if/how DC axons are activated by SCS.

### Activation of DC axons

To test if SCS activates DC axons, we recorded from the L4 dorsal root ganglion (DRG) to detect antidromically propagated spikes. We isolated single-unit activity by subtracting the stimulus artifact and ECAPs recorded on multiple contacts of a multielectrode array (MEA) from the signal recorded exclusively on one contact, presumably closest to the unit (**Supplementary Fig. 2**). SCS-responsive units were found by adjusting the MEA position during 2 Hz “search” SCS. All units identified during 2 Hz search (labeled 2Hz+ units) responded vigorously when subsequently tested with c-SCS (blue) or kf-SCS (red) (**Fig. 2A**). Most units fired at 50 spk/s during c-SCS by responding to each pulse with one spike, whereas many units responded to kf-SCS with firing rates >50 spk/s (**Fig. 2B**). Median firing rates for 2Hz+ units during c-SCS (50 spk/s) and kf-SCS (47.3 spk/s) were not significantly different (*U*=107, *p*=0.45, Mann-Whitney test) but, because of the skewed distribution, average firing rate was 60% higher during kf-SCS than during c-SCS (63.8 vs 40.0 spk/s); here, firing rate is calculated for each neuron based on its spike count over the first 20 s of SCS. Additional units not identified during 2 Hz search (labeled 2Hz-units) were observed almost exclusively during kf-SCS (χ^2^=4.68, *p*=0.03) and fired at a lower rate (4.6 spk/s) than 2Hz+ units (*U*=10.0, *p*=0.001, Mann-Whitney test). The receptive field (RF) was identified in nearly all 2Hz-units but in only a minority of 2Hz+ units (χ^2^=8.81, *p*=0.003). Because they did not respond to isolated (2 Hz) SCS pulses, the conduction velocity of 2Hz-units was not formally measured but their responsiveness to tactile stimulation identified them as low-threshold mechanoreceptive afferents. Amongst 2Hz+ units, conduction velocity did not differ significantly between those with or without an identified RF (*T*_31_=-0.047, *p*=0.96, unpaired t-test) (**Fig. 2C**).

**Figure 2.**
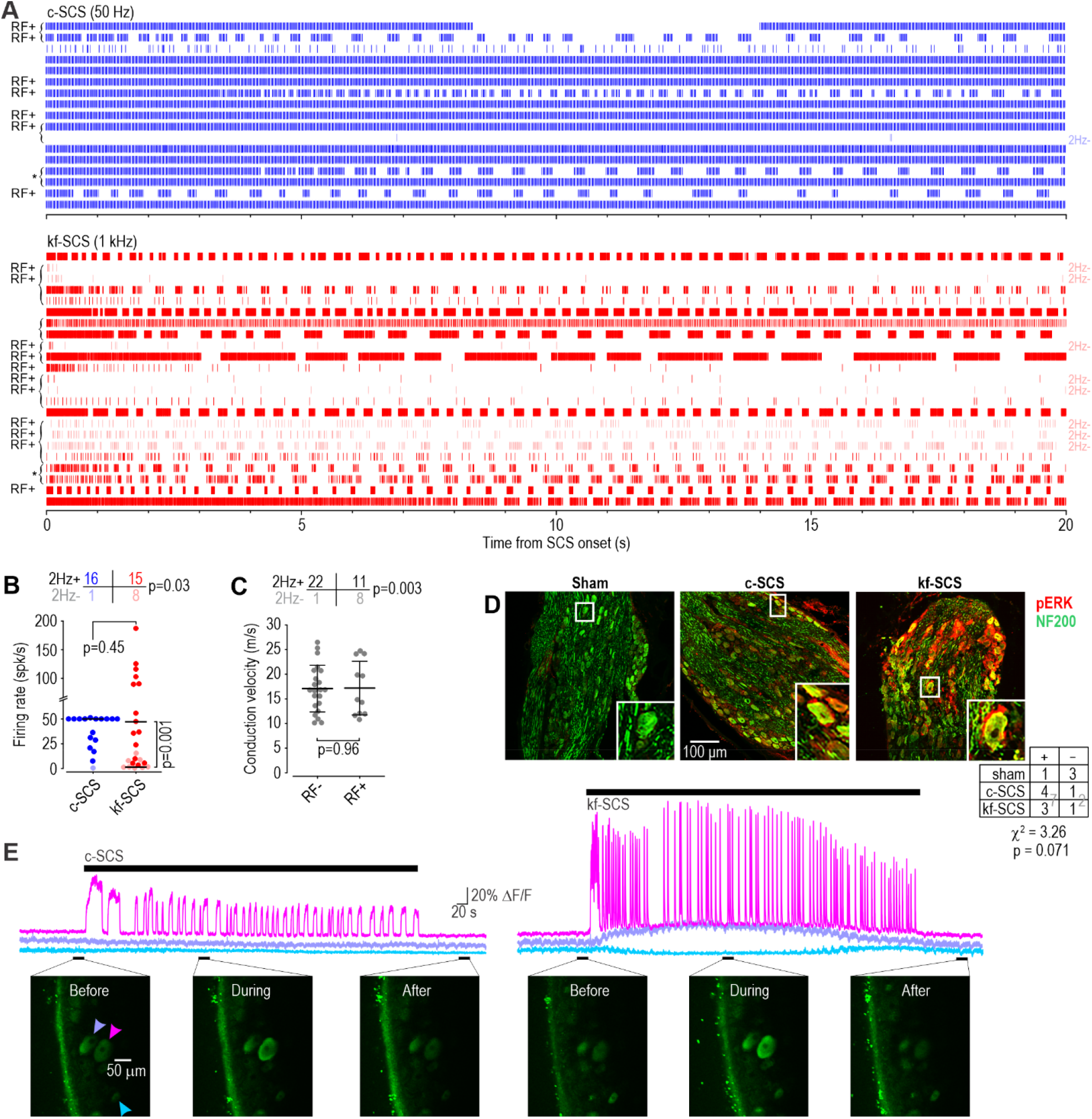
Both forms of SCS activate DC axons. **(A)** Rasters depict each spike as a tick. Intermittent gaps in spiking reflect bursts. Each row is a different unit. *RF+*, unit with identified receptive field. *2Hz-* (pale), unit not activated during 2 Hz search stimulation. Brackets, simultaneously recorded units. *, units analyzed in Fig 3E,F. **(B)** Amongst 2Hz+ units (dark), the median firing rate (based on spike count over the first 20 s of SCS) did not differ between c-SCS and kf-SCS (*U*=107, *p*=0.45, Mann-Whitney test) yet mean rate during kf-SCS (63.8 spk/s) was higher than during c-SCS (40.3 spk/s) because of the skewed distributions. 2Hz-units (pale) were observed almost exclusively during kf-SCS (χ^2^=4.68, *p*=0.03) and fired at a lower rate (4.6 spk/s, *U*=10.0, *p*=0.001, Mann-Whitney test). Bars show median. **(C)** The receptive field (RF) was identified in more 2Hz-units than in 2Hz+ units (χ^2^=8.81, *p*=0.003). Conduction velocity (based on latency of spikes conducted antidromically from the stimulation site to the DRG recording site) did not differ between RF+ and RF-units (*T*_31_=-0.047, *p*=0.96; unpaired T-test). Bars show mean±SD. **(D)** pERK (red) was observed in myelinated neurons identified by neurofilament 200 (NF200) expression (green) after c-SCS and kf-SCS, but not after sham. Table summarizes number of animals scored as pERK+ or – by a blinded evaluator (χ^2^=3.26, *p*=0.071 using data pooled across SCS conditions). **(E)** Two-photon GCaMP imaging. Example images show the same field of view imaged during c-SCS (left) and kf-SCS (right). Arrowheads point to three cell bodies whose fluorescence changes (∆F/F) are plotted against time. Movement artifacts during kf-SCS caused slow fluorescence changes distinct from SCS-evoked activity (pink) whose intermittent fluctuations are consistent with bursting. See Supplementary Fig. 3 for additional data and analysis.

Since activation of DC axons by kf-SCS is inconsistent with past work^15, 16^ and since recordings can be compromised by kf-SCS^19, 30^, we verified our electrophysiological results using two additional methods, namely immunostaining for the activity marker phosphorylated extracellular signal-related kinase^31^ (pERK; **Fig. 2D**) and two-photon calcium imaging (**Fig. 2E**; see also **Supplementary Fig. 3**). Both methods confirmed that c-SCS and kf-SCS applied at 50% MT selectively activate large myelinated afferents.

### Differentially synchronized spiking in activated axons

The main difference between c-SCS and kf-SCS is evident in the pattern of evoked spiking. During c-SCS (**Fig. 3A**), each SCS pulse typically evoked one spike (1:1 entrainment), meaning nearly all spikes occurred with an interspike interval (ISI) of 20 ms (=1/50 Hz). Population firing rate jumped abruptly after each pulse, when most neurons fired a spike, then dropped back to zero until the next pulse; here, firing rate is calculated from all recorded neurons based on the number of spikes per 2 ms bin. During kf-SCS (**Fig. 3B**,**C**), spikes phase locked to SCS pulses but occurred on only a subset of pulses, producing ISIs at *multiples* of the 1 ms (=1/1 kHz) interpulse interval. ISIs as short as 2 ms (1:2 entrainment; 1 spike on every 2 pulses) were observed but most ISIs were between 3 and 6 ms (1:3 to 1:6 entrainment). Long (>50 ms) ISIs correspond to the gaps between bursts (see rasters on Fig. 2B). This pattern emerges because units skip (i.e. fail to respond to) pulses arriving while the unit is still refractory from the previous spike (see below). By skipping a variable number of pulses, different units tend to fire on different pulses, desynchronizing the population response, as evident from the blunted fluctuations in population firing rate (**Fig. 3B top**). **Figure 3D** summarizes the cumulative probability distribution of ISIs. These patterns of spiking – synchronous during c-SCS vs asynchronous during kf-SCS – persisted for the duration of testing (**Supplementary Fig. 4**).

**Figure 3.**
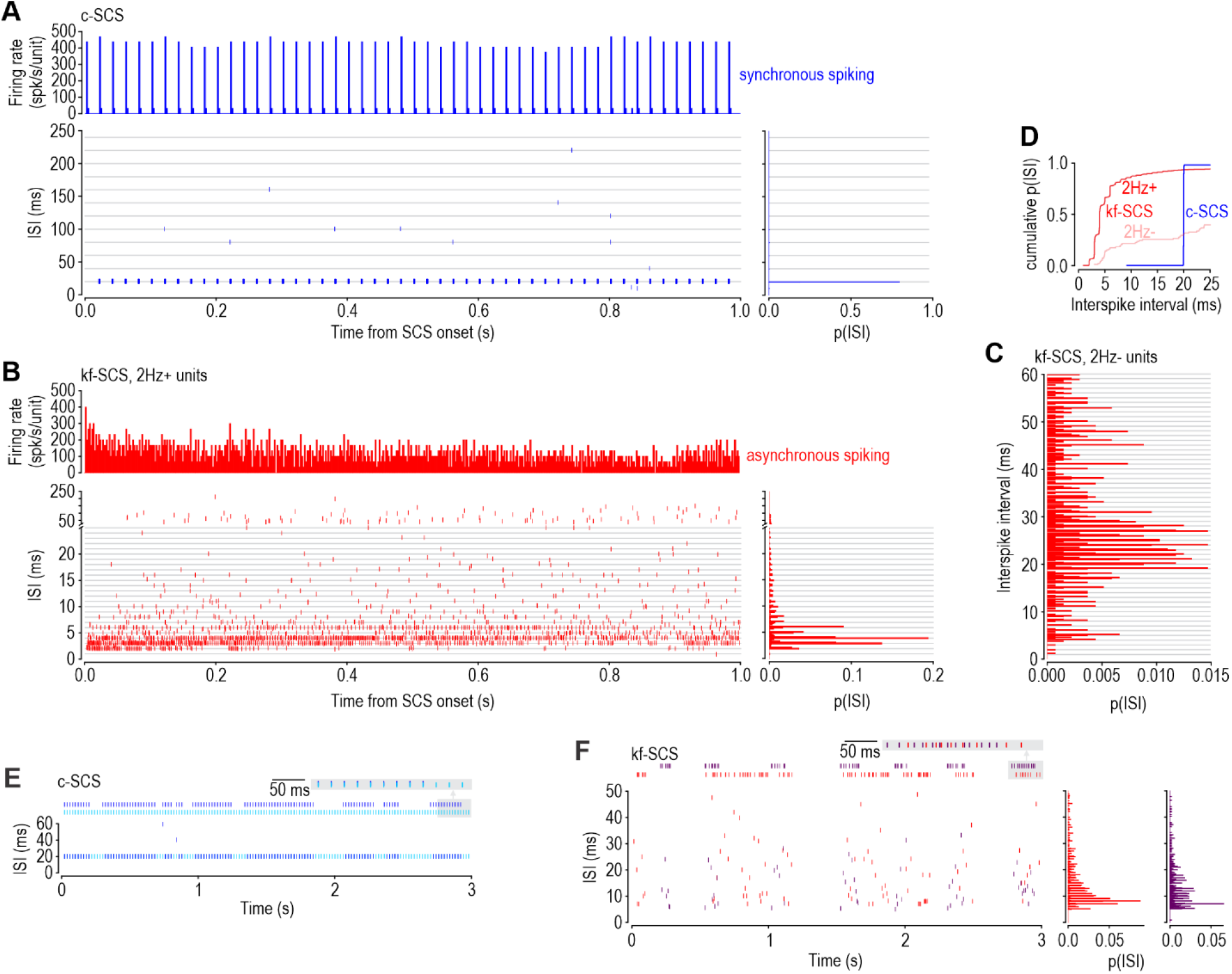
Spiking evoked by kf-SCS is less synchronized than c-SCS-evoked spiking. Gray lines represent integer multiples of the interpulse interval: 20 ms for c-SCS vs. 1 ms for kf-SCS. **(A)** During c-SCS, nearly all spikes (from 17 units) occurred at 20 ms intervals because of 1:1 entrainment with SCS pulses. Population firing rate (calculated using a 2 ms-wide bin) fluctuates because most neurons fire synchronously after each SCS pulse and then remain quiescent until the next pulse. **(B**,**C)** During kf-SCS, spikes (from 23 units) occurred at various multiples of 1 ms. Most ISIs were between 3 and 6 ms in 2Hz+ units **(B)** whereas longer ISIs occurred in 2Hz-units **(C)**. This pattern is consistent with irregular skipping (see Figure 4), which blunts fluctuations in the population firing rate by causing neurons to fire *asynchronously* (i.e. on different pulses). **(D)** Cumulative probability distribution of all ISIs analyzed in A-C. **(E**,**F)** Responses from two simultaneously recorded units (marked with * on Figure 2A). Both units fired on most pulses (i.e. synchronously) during c-SCS **(E)** but on different pulses (i.e. asynchronously) during kf-SCS **(F)**, despite having similar intraburst ISI distributions.

Data in Figure 3A-D were collected from several rats and aligned based on stimulus timing. But neurons might skip the *same* SCS pulses and continue to spike synchronously during kf-SCS if they experienced correlated noise^32^; testing this requires simultaneously recorded units. Though MEAs enable multi-neuron recordings, we rarely recorded more than one 2Hz+ unit simultaneously (see brackets on Fig. 2A) during low-amplitude SCS, consistent with the sparse activation suggested by calcium imaging (see Supplementary Fig. 3) and with past work^33^. As illustrated in a pair of 2Hz+ units recorded simultaneously during c-SCS and kf-SCS, both units fired on the same pulses during c-SCS (**Fig. 3E**) but on different pulses during kf-SCS (**Fig. 3F**). In this and all other pairs (including all 2Hz-units, which were always recorded with a 2Hz+ unit because of our search protocol), bursts sometimes occurred concurrently but spikes never synchronized (**Supplementary Fig. 5**). These results show that noise is uncorrelated and, therefore, encourages desynchronized spiking during kf-SCS.

Irregular skipping accounts for asynchronous spiking during kf-SCS (see below) with other potential mechanisms such as temporal dispersion making little contribution (**Supplementary Fig. 6**). Asynchronous spiking was also observed during 0.5 kHz and 5 kHz SCS (**Supplementary Fig. 7**). We used the same pulse waveform for all SCS frequencies but c-SCS is usually applied with a longer pulse, and longer pulses are typically applied at lower amplitude, which might be less synchronizing. When c-SCS was re-tested using 200 µs-long pulses, MT was predictably lower (see Supplementary Fig. 1) but c-SCS applied at 50% of this lower MT still evoked synchronous spikes (**Supplementary Fig. 8**). The differentially synchronized spiking evoked by c-SCS and kf-SCS prompted us to investigate the basis for and consequences of spike (de)synchronization.

### Basis for differentially synchronized spiking

We hypothesized that skipping occurs when the interpulse interval is shorter than the axon refractory period. We tested this by modeling spike generation as a stochastic process (**Fig. 4A**). A random number is chosen each time an SCS pulse occurs. If the number exceeds threshold, a spike is generated (*i*), threshold is incremented (*ii*), and the distribution from which future numbers are drawn shifts according to time-dependent changes in post-spike excitability (*iii*). The refractory period decreases the probability of a spike occurring at short (<3 ms) intervals but the subsequent increase in excitability (i.e. superexcitability^34^) increases the probability of a spike occurring at intermediate (3-10 ms) intervals. At the same time, threshold decays slowly back to baseline. Bursts arise through competition between mechanisms *ii* and *iii*: a burst terminates when superexcitability (after each spike) fails to offset the threshold increase (accumulated across multiple spikes); the next burst starts only after the increase in threshold has worn off. Stimulating (i.e. picking random numbers) at 1 ms intervals to model kf-SCS capitalized on fast fluctuations in post-spike excitability to produce an intraburst ISI distribution like in kf-SCS experiments (**Fig. 4B**, cf. Fig. 3A). Stimulating the same models at 20 ms intervals to model c-SCS produced spikes at regular 20 ms intervals (**Fig. 4C**). The key difference is indeed whether the interpulse interval is shorter or longer than the axon’s refractory period.

**Figure 4.**
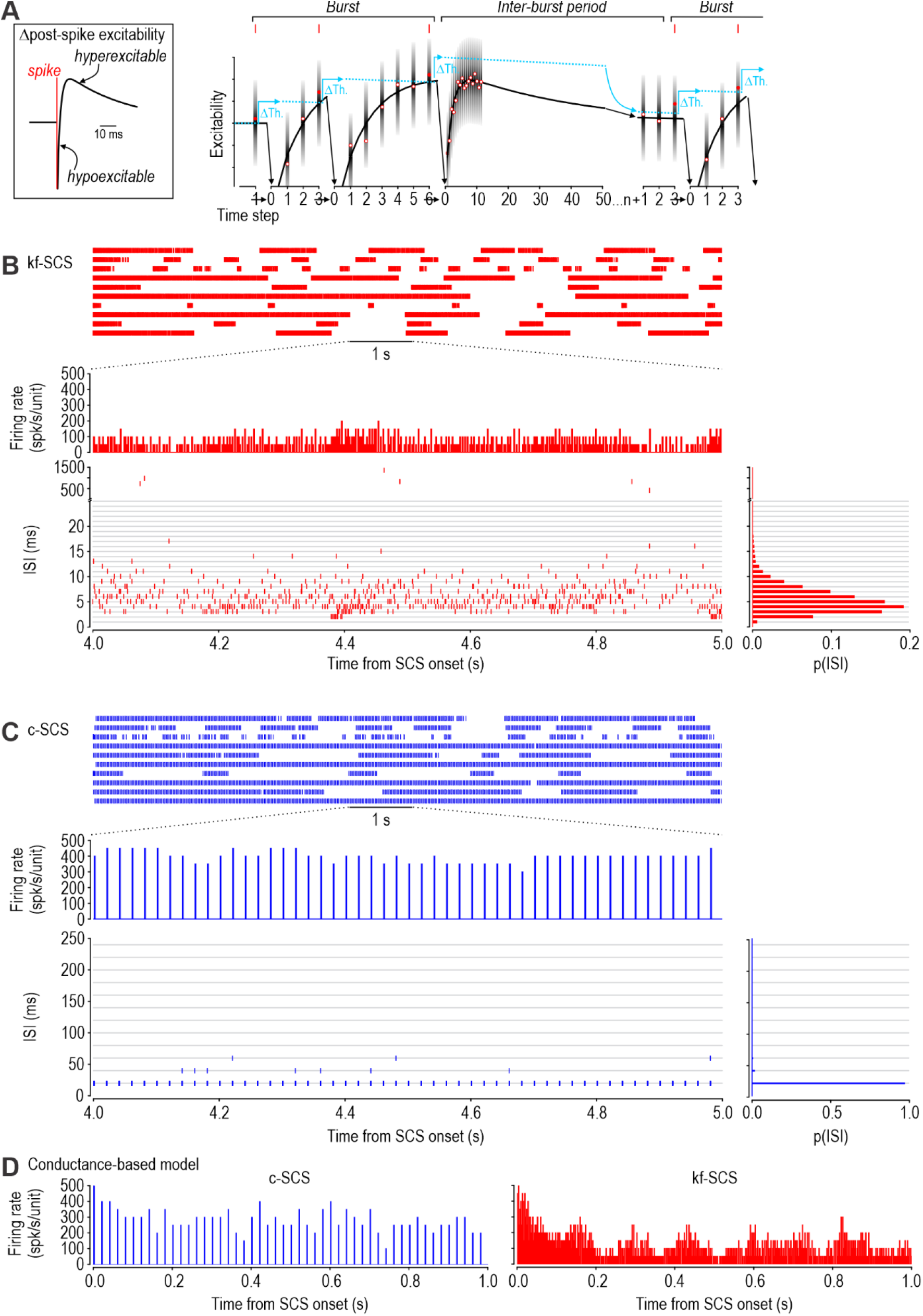
Irregular skipping occurs when an axon is stimulated during its refractory period. **(A)** Phenomenological model. At each time step, a random number (red circle) is drawn from a Gaussian distribution (grayscale shading) to model spike generation as a stochastic process. If the number exceeds threshold (dotted cyan line), a spike occurs and threshold is stepped up (Δ*Th*) before slowly relaxing back toward baseline. On a faster timescale, the Gaussian distribution shifts according to changes in post-spike excitability (inset on left). Fast excitability changes dictate the intra-burst spiking pattern while slow changes in threshold dictate burst duration (see Results). **(B)** During kf-SCS (random numbers drawn at 1 ms intervals), model axons responded with asynchronous spiking. Parameter values differed slightly across the ten models to introduce heterogeneity and random numbers were chosen independently for each model to simulate uncorrelated noise. The resulting ISI distribution and firing rate histogram accurately reproduced experimental data in Figure 3A. **(C)** The same models (from B) spiked synchronously during c-SCS (random numbers drawn at 20 ms intervals). **(D)** Conductance-based axon models likewise exhibited asynchronous and synchronous firing during c-SCS and kf-SCS, respectively. See Supplementary Figs. 9 and 10 for additional analysis of phenomenological models and Supplementary Fig. 11 for additional analysis of conductance-based models.

Simulating the same model axons with different noise confirmed that uncorrelated noise and cellular heterogeneity are each sufficient to desynchronize spiking evoked by kf-SCS, yet neither desynchronizes the spiking evoked by c-SCS (**Supplementary Figure 9**). Gilbert et al. recently showed that pulses with perithreshold amplitude evoke spikes intermittently, causing desynchronization even at low frequencies^35^. Indeed, some perithreshold pulses can be rendered subthreshold by noise alone, without requiring spike-induced fluctuations in excitability (although such fluctuations expand the pulse amplitude range over which desynchronization occurs, abrogating the need to titrate pulse amplitude). If axons experience uncorrelated noise and/or their refractoriness wanes at different rates (especially considering that each axon experiences SCS differently depending on its diameter and location in the electric field^36, 37^), pulses arriving while axons are still refractory will cause different axons to respond to different pulses, thus desynchronizing the population response – we refer to this as overdrive desynchronization. Suprathreshold pulses occurring at longer intervals do not reveal this underlying heterogeneity, allowing spikes to entrain reliably to suprathreshold pulses and thus remain synchronous. Based on these insights and a typical refractory period of ∼3 ms, we predicted and confirmed (see **Supplementary Fig. 10**) that overdrive desynchronization should start at SCS frequencies near 333 Hz (=1/3 ms). We also confirmed that overdrive desynchronization occurs in a conductance-based axon model during kf-SCS, but not during c-SCS (**Fig. 4D** and **Supplementary Fig. 11**). These results explain why axons respond so differently to different SCS frequencies.

### Consequences of differentially synchronized spiking

Synchronous spikes are liable to evoke sensation (paresthesia) because their transmission through downstream circuits is less readily blocked by feedforward inhibition (**Supplementary Figure 12**). But whereas synchronous DC axon spiking is necessary for paresthesia – consistent with the correlation between ECAP amplitude and paresthesia intensity (see below) – asynchronous spiking is sufficient to produce analgesia. Inhibitory neurons in the spinal dorsal horn are activated during kf-SCS^21, 22^, which, according to our results, occurs via synaptic input from activated DC axons. Synaptic input need *not* be synchronized since inhibitory neurons behave as integrators that summate temporally dispersed inputs^38^; indeed, they are naturally activated by input from slow-adapting mechanoreceptive afferents that spike asynchronously during vibrotactile stimulation^39^. Desynchronized activation of DC axons thus avoids paresthesia without compromising analgesia.

Next, we sought a surrogate marker for overdrive desynchronization. In rat experiments, the amplitude of ECAPs recorded from inside the DRG correlated with pulse amplitude during c-SCS (**Fig. 5A left**) because, unlike all-or-none single-unit spikes (**Fig. 5A, right**), the ECAP is a multi-unit phenomenon whose amplitude depends on the number of *synchronously* activated units^40, 41^. Synchrony is key. If units fire at the same rate during c-SCS and kf-SCS but distribute their spikes evenly over 20× more pulses in the latter case, then 95% fewer spikes occur per pulse during kf-SCS, meaning ECAPs should be 95% smaller. More realistically, ECAP amplitude is liable to fluctuate during kf-SCS, depending on how spikes distribute across pulses, but will invariably be smaller than during c-SCS. As predicted, ECAPs were large during c-SCS but were small and irregular during kf-SCS (**Fig. 5B**). Large voltage fluctuations were evident only at the onset of kf-SCS (bottom trace in Fig. 5B), consistent with desynchronization arising gradually through the cumulative effect of irregular skipping^42^ (see also **Supplementary Fig. 11**)

**Figure 5.**
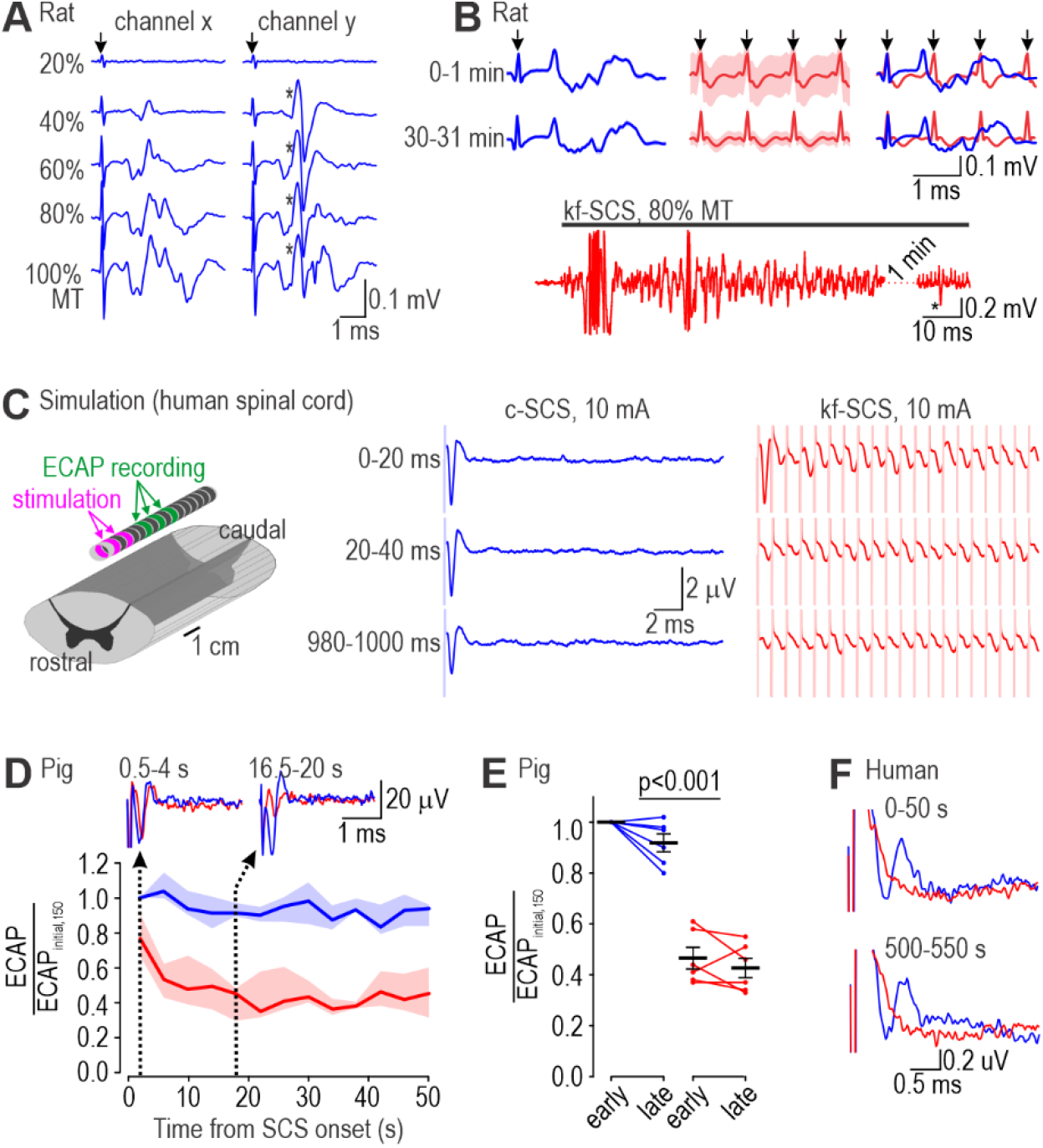
ECAPs are prominent only during c-SCS. **(A)** Simultaneous recordings in rat DRG from two electrodes during c-SCS. ECAP recorded on channel x grew steadily as pulse amplitude was increased from 20% to 100% MT whereas a full-sized single-unit spike (*) appeared on channel y at 40% MT, partly obscuring the ECAP. Arrows mark stimulus artifacts. Beyond showing that ECAP amplitude scales with pulse amplitude, these data highlight that single-units spikes can occur in the absence of large ECAPs. **(B)** Average ECAP (±standard deviation, shading) over first and 30^th^ minute of c-SCS (blue) and kf-SCS (red) tested in the same rat at 80% MT. Right panel shows overlay. A consistent ECAP was evident only during c-SCS. Irregular fluctuations during kf-SCS averaged out, leaving only stimulus artifacts; standard deviation was large in the first minute, before fluctuations subsided. Bottom trace shows large, irregular voltage fluctuations at the onset of kf-SCS; trace is interrupted for 1 min for comparison with later response, where single-units spikes (*) are evident in the absence of ECAPs (see also Supplementary Fig. 2). ECAPs in A and B were recorded from inside the DRG and thus differ in shape and amplitude from ECAPs recorded epidurally in C-F. **(C)** ECAP simulations in human spinal cord model. Schematic shows spinal cord (whose dorsal column was populated with conductance-based axon models) and electrodes for stimulation and recording. Simulated ECAPs are shown for three 20 ms-long epochs; artifacts are truncated and shaded for clarity. Whereas c-SCS evoked large ECAPs that persisted throughout the 1 s-long simulation, a recognizable ECAP was evident only at the onset of kf-SCS before giving way to small, irregular voltage fluctuations. **(D)** ECAPs were measured in 6 pigs during SCS at 150 Hz (c_150_-SCS) and during kf-SCS. Averaging every 4 consecutive trials revealed that ECAPs are initially large during kf-SCS but rapidly attenuate. Line and shading show median and interquartile range, respectively. Sample ECAPs from one pig are shown at two time points. Data from each pig were normalized by the average early ECAP during c_150_-SCS. **(E)** ECAP amplitude differed significantly between SCS protocols (*F*_1,5_=141.97, *p*<0.001, two-way repeated measures ANOVA) but not between early and late epochs (*F*_1,5_=4.15, *p*<0.10) nor was there any interaction (*F*_1,5_=1.20, *p*<0.23). Each data point is from one pig and represents the ECAP averaged over the first or last 50 s of 500 seconds of SCS (“early” and “late”, respectively). Data were normalized like in D and summarized as mean±SEM. Post-hoc SNK tests are reported on graph. **(F)** Sample ECAPs from a chronic pain patient. With pulse amplitude set to just above paresthesia threshold during c_150_-SCS, ECAPs were evident during c_150_-SCS but not during kf-SCS, consistent with presence or absence, respectively, of reported paresthesia. Given the consistent pulse amplitude and width, 6.67× more charge was delivered during kf-SCS, yet there was no paresthesia.

To verify the effects of desynchronization on ECAPs, conductance-based human axon models were incorporated into a human spinal cord model and stimulated by electric field changes, and their spiking was used to construct ECAPs (see Methods). As predicted, the synchronous spiking evoked by c-SCS produced consistently large ECAPs whereas the asynchronous spiking evoked by kf-SCS produced much smaller ECAPs (**Fig. 5C**). Equivalent experiments were conducted in six pigs. Conventional-SCS was tested at 150 Hz (henceforth referred to as c_150_-SCS) because the active recharge required for short pulses was unavailable for frequencies <150 Hz on the stimulators used. To measure ECAPs uncorrupted by stimulus artifacts during kf-SCS, stimulation was paused for 6.67 ms after 500 ms-long stimulus blocks to give an artifact-free interval equivalent to the interpulse interval for c_150_-SCS (1/150 Hz = 6.67 ms); the ECAP was measured during these pauses in kf-SCS and at the equivalent interval during c_150_-SCS, namely every 76^th^ pulse, so that the number of averaged traces was equivalent between conditions. Like in simulations, ECAP amplitude remained stable during c_150_-SCS but dropped rapidly after the initial phase of kf-SCS (**Fig. 5D**). Analyzed over a longer period, kf-SCS produced significantly smaller ECAPs than c_150_-SCS (*Q*=16.850, *p*<0.001; SNK test) despite, for each pig, using the same pulse amplitude and width (i.e. charge per pulse) for c_150_-SCS and kf-SCS. Lastly, the same protocol was carried out in a chronic pain patient. Pulse amplitude was adjusted to slightly above perception threshold for c_150_-SCS, thus causing paresthesia, but when SCS frequency was increased to 1 kHz without changing other parameters, paresthesia was absent, consistent with negligible ECAPs during kf-SCS (**Fig. 5F**). Given other work showing that paresthesia intensity correlates with ECAP amplitude when pulse amplitude is varied^43^, our results show that desynchronizing DC spiking (by increasing SCS frequency) is an alternate way to minimize ECAPs and their associated paresthesia. Unlike reducing pulse amplitude, increasing SCS frequency reduces paresthesia without compromising analgesia – by reducing synchrony without reducing firing rate. In other words, a high rate of asynchronous spiking in DC axons explains paresthesia-free analgesia.

## DISCUSSION

Our results show that kf-SCS and c-SCS both activate DC axons (Fig. 2) but that kf-SCS activates those axons less synchronously than c-SCS (Fig. 3). Desynchronization occurs because, when pulses occur at an interval shorter than the axon refractory period, axons fire on only a subset of pulses, and different axons fire on different pulses because of uncorrelated noise and cellular heterogeneity (Fig. 4). Desynchronization leads to smaller ECAPs (Fig. 5). Given past work showing a correlation between ECAP amplitude and paresthesia intensity (see below), desynchronization of the population response explains how DC axons can fire at high rates without producing paresthesia. Our results do not address how DC axon activity mediates analgesia, but given that c-SCS and kf-SCS produce comparable modulatory effects (Fig. 1), their mechanisms are probably more similar than widely speculated. If charge (dose) is correlated with analgesia^44^ but limited by paresthesia^10^, then applying pulses at higher rate – at the same or even lower amplitude – can avoid paresthesia while maximizing analgesia (**Fig. 6**).

**Figure 6.**
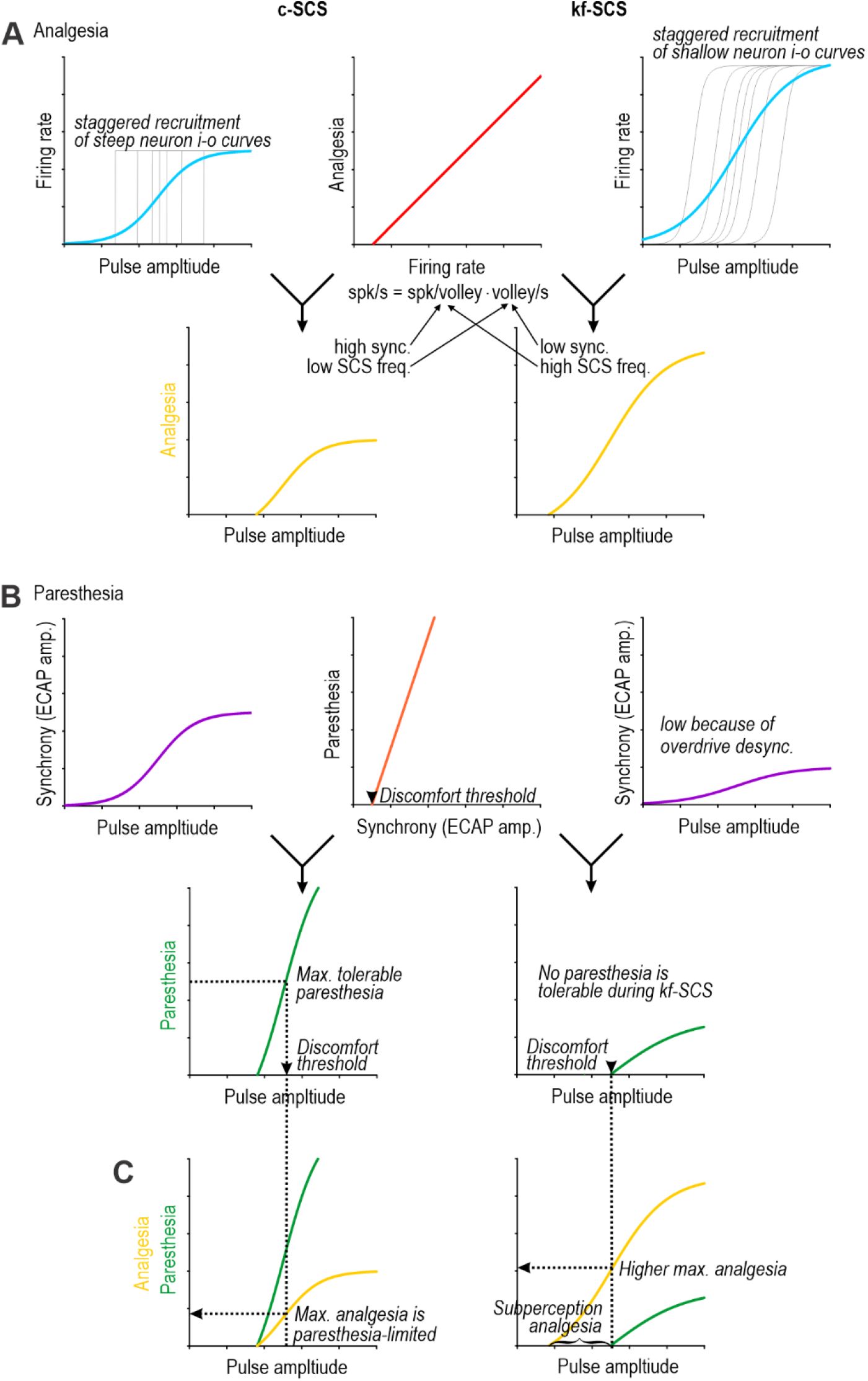
Mathematical model explaining the dependence of analgesia and paresthesia on the rate and synchrony of SCS-evoked spiking. **(A)** Analgesia depends on firing rate (red curve), which depends on pulse amplitude (blue curve) and SCS frequency (cf. blue curves on left and right). Convolving the red and blue curves yields the gold curve relating analgesia to pulse amplitude. **(B)** Paresthesia depends on spike synchrony (orange curve), which depends on pulse amplitude (purple curve) and SCS frequency (cf. purple curves on left and right). Convolving the orange and purple curves yields the green curve relating paresthesia to pulse amplitude. Gold and green curves differ between c-SCS (left) and kf-SCS (right) entirely because of differences in the blue and purple curves that reflect differences in the rate and pattern of SCS-evoked spiking; overdrive desynchronization during kf-SCS ensures low synchrony. Importantly, low synchrony combined with high SCS frequency (during kf-SCS) can evoke the same (or even greater) firing as high synchrony combined with low SCS frequency (during c-SCS). Blue curves differ between c-SCS and kf-SCS because of how axons are recruited, as illustrated by thins lines depicting input-output (i-o) curves for individual axons: axons switch abruptly from quiescence to firing at 50 spk/s when their threshold is reached during c-SCS, but are recruited more gradually and reach higher firing rates during kf-SCS. **(C)** Overlaying the gold and green curves reveals the relationship between analgesia and paresthesia. Unless pulse amplitude, pulse width, and rate are carefully titrated with respect to each other, no analgesia occurs without paresthesia during c-SCS (left), unlike during kf-SCS (right). Uncomfortable paresthesia limits analgesia in both cases, but by activating DC axons more strongly (vertically expanding the gold curve) and minimizing their synchronization (right shifting the green curve), kf-SCS can produce greater analgesia.

The intrinsic excitability of DC axons defines the stimulus conditions producing overdrive desynchronization. Axons skip (i.e. fail to respond to) pulses delivered at intervals shorter than their refractory period. Skipping has been reported previously, for example, in motor axons during electrical stimulation^45^, in sensory axons during vibrotactile stimulation^46^, and in simulations^47^. The relative refractory period estimated here to be ∼3 ms is consistent with threshold tracking measurements using paired pulses^34^. That value identifies 333 Hz (=1/3 ms) as the SCS frequency at which overdrive desynchronization starts, although, in reality, it develops gradually around that frequency. The refractory period gives way to supernormal excitability resulting from discharge of current that builds up on the internode during a spike^48^. This phase, which corresponds intracellularly with an afterdepolarization (ADP), interacts with slower changes in excitability to produce bursting (which may induce analgesia differently than the mostly tonic spiking evoked by c-SCS). Temporal summation of subthreshold depolarization (evoked by low amplitude pulses) involves the same process underlying the ADP, and is maximal for an interpulse interval of ∼5 ms^49, 50^. This explains why 2Hz-units were activated almost exclusively during kf-SCS: individual pulses were subthreshold (unlike for 2Hz+ units) but their depolarizing effect could summate across 1 ms intervals (during kf-SCS) but not across 20 ms intervals (during c-SCS). During c-SCS, an axon transitions abruptly from quiescence to firing at 50 spk/s when pulse intensity reaches threshold. During kf-SCS, the same axon responds to subthreshold pulses with low-rate firing (thanks to summation) and gradually increases its firing rate as pulse amplitude is increased (and fewer pulses are skipped). Activation of more DC axons (thanks to summation of subthreshold pulses) to a firing rate higher than the 50 spk/s possible with 1:1 entrainment during c-SCS enables greater analgesia for a given pulse amplitude during kf-SCS (see Fig. 6). This assumes pulse width is constant across SCS frequencies, which is often not the case; charge per pulse (= pulse amplitude × width) is more relevant than pulse amplitude per se. The bursting and temporal summation associated with kf-SCS may also produce effects absent during c-SCS, and require further investigation.

Spike synchrony is important for sensation^51^ and, more generally, for transmission of neural information^52, 53^. Sensations of flutter and vibration normally arise from vibrotactile stimuli < or >50 Hz, respectively, that evoke precisely timed (synchronous) spikes in mechanoreceptive afferents^54, 55^ (consistent with changes in paresthesia quality when SCS frequency is varied across that range^56^). Perception arguably depend more on spiking pattern than on which type of afferent is activated^57^, but differences in mechanotransducer tuning normally result in differential activation^58^, complicating disambiguation. How we perceive frequency (as pitch) and intensity (as loudness) has been debated for nearly a century in the auditory system^59^. For *tonic* electrical stimulation in the somatosensory system, Graczyk et al.^60^ showed that “loudness” (i.e. perceived touch intensity) depends jointly on pulse rate and pulse charge, which can be summarized as charge rate (= charge/pulse × pulse/sec). For *bursting* electrical stimulation, Menia and Van Doren^61^ showed that “loudness” and “pitch” (i.e. perceived touch quality) correlate respectively with burst charge and burst rate. In the latter study, bursts comprised high-rate pulses; participants perceived the burst rate (2-64 Hz) rather than the intraburst pulse rate (1 kHz). Both studies are relevant for understanding SCS-evoked paresthesia (see below).

The intensity (loudness) of paresthesia correlates with ECAP amplitude when pulse amplitude is varied, with sensory threshold (ST) coinciding roughly with ECAP threshold (i.e. the minimum charge per pulse required to produce a detectable ECAP)^43, 62, 63^. If a spike from a single axon produces a ∼1 µV signal recorded epidurally, then several dozen active units are required to produce a detectable ECAP^62, 64^, consistent with evidence that up to 20% of fibers can spike without generating a detectable ECAP^65^. Gmel et al. showed that increasing SCS frequency (as high as 455 Hz in one subject) without changing other stimulus parameters produced progressively smaller ECAPs but more intense (and uncomfortable) paresthesia^63^; the reported changes in paresthesia arguably reflect a change in pitch (and sensitivity to different pitches) rather than a pure increase in loudness (see above). The diminishing ECAP amplitude with increasing SCS frequency Gmel et al. reported is consistent with our results, but the human participant in our study no longer felt any paresthesia – and ECAPs disappeared – when SCS frequency was increased from 150 Hz to 1 kHz, despite applying the same charge per pulse; normally (i.e. clinically), the charge per pulse is reduced as SCS frequency is increased, which further reduces the likelihood of paresthesia. Recent work showed that low-frequency (<100 Hz), small-amplitude pulses produce pain relief without paresthesia^66^ by activating DC axons in a sparse, irregular pattern which engages spinal inhibition^35^. This argues that high-rate and/or low-amplitude SCS achieves paresthesia-free pain relief through asynchronous activation of DC axons.

Classic microstimulation experiments showing that activation of a single somatosensory afferent can be perceived^67-69^ seem to contradict our claim that several axons must be synchronously activated to produce paresthesia. Notwithstanding technical concerns^70 71^, perception of activity in a single afferent likely requires absence of activity in other afferents because of surround suppression; specifically, stimulation outside a neuron’s receptive field (or, equivalently, coincident activation of neurons with non-overlapping receptive fields, as occurs during SCS) tends to inhibit that neuron or its downstream targets^72, 73^. When many afferents are co-activated (by tactile stimulation or by SCS), huge amounts of feedforward, lateral, and feedback inhibition are engaged, dampening the impact of input from any one afferent. Neurons projecting from DC nuclei to thalamus experience strong surround inhibition activated by primary afferent input^74^. Surround Inhibition also dampens pain signals (see Introduction) as well as evoked potentials and reflexes (see Fig. 1). The intermittent, highly synchronized spike volleys underlying paresthesia overcome this inhibition by arriving moments before the onset of feedforward inhibition and 10s of milliseconds after inhibition engaged by the preceding volley, when inhibition is at its weakest (see Supplementary Fig. 12). In contrast, continuous desynchronized spike volleys produce gap-free inhibition and evoke excitatory input too small to overcome that inhibition.

By uncovering the intrinsic properties that dictate how an axon responds to pulse trains and how individual axon responses combine to produce the population response, our results reveal how to sculpt the population response using strategically chosen stimulation parameters. Whereas reducing paresthesia by reducing pulse amplitude reduces analgesia^75^, overdrive desynchronization allows the use of high-amplitude pulses. That said, since charge rate (charge/sec = charge/pulse × pulse/sec) is correlated with pain relief^44^, pulse amplitude can be reduced as pulse rate is increased, although pulse amplitude reduction is limited by pulse width, which must remain short given the short interpulse intervals required for kf-SCS. Stimulating at high rate with the widest possible pulse (high duty cycle) in order to apply the lowest effective pulse amplitude amounts to high-density SCS^76^. Such parameter combinations drive DC axons strongly but asynchronously, which conveys good analgesia without paresthesia, but consumes significant energy. Effective, energy efficient SCS can be achieved by co-adjusting rate, pulse amplitude, and pulse width^77, 78^.

To conclude, our results show that DC axons are activated synchronously during c-SCS and asynchronously during kf-SCS. Desynchronization of spiking attenuates ECAP amplitudes. Given the strong correlation between ECAP amplitude and paresthesia intensity, asynchronous activation of DC fibers explains pain relief by paresthesia-free SCS. Overdrive desynchronization explains how this occurs during kf-SCS.

## METHODS

### Rat experiments

All experimental procedures in rats were approved by the Animal Care Committee at the Hospital for Sick Children (protocol #47072). Adult Sprague Dawley rats (200-350 g) were obtained from Charles River, Montreal. Males were used for most experiments but calcium imaging was conducted in both sexes; no sex differences were observed.

#### Spinal cord stimulation

Healthy rats were anesthetized by inhalation of 4% isoflurane followed by urethane (1.2-1.5 g/kg i.p.). To implant the SCS lead in the epidural space, the T13 vertebra was removed. A 4-contact preclinical lead (Boston Scientific Neuromodulation, Valencia, CA) was implanted midline with the tip beneath the T11 vertebra (Fig. 1A), which corresponds to the T13 spinal level^79^. The two distal contacts (1 mm separation) were used to deliver constant-current stimulation using a narrow bipole (to preferentially activate DCs rather than dorsal roots^80^). Stimulation was controlled by a therapy-grade Precision Spectra™ External Trial Stimulator from Boston Scientific^81^ with non-commercial firmware change that enabled SCS from 2 Hz to 5 kHz using a symmetrically biphasic square wave (40 µs per phase). Except in a subset of experiment where a longer biphasic pulse was tested at 50 Hz (since c-SCS is usually applied using a longer waveform, e.g. 200 µs of active charge followed by passive recharge), the same short waveform was used across SCS frequencies to enable back-to-back comparison. Amplitude was set to 50% of MT or was adjusted to between 40-60% of MT while searching for units (with 2 Hz SCS). To determine MT, the amplitude of SCS pulses (applied at 2 Hz) was increased until muscle twitches were observed.

It should be noted that mechanoreceptor and proprioceptor axons are differentially positioned in the DC, which has implications for their activation by SCS. Electrophysiological analysis in cats showed that mechanoreceptor axons are more superficial than proprioceptor axons^82^. Molecular genetic analysis in mice showed that mechanoreceptor axons are more medial than proprioceptor axons^83^. Assuming no species differences, this positioning favors activation of mechanoreceptor axons given the midline position of the SCS lead whereas the larger caliber of proprioceptor axons favors their activation^84^.

#### Dorsal root ganglia (DRG) recordings

For DRG recordings, a laminectomy was performed to expose the L4 DRG. Axons forming the sciatic nerve originate predominantly from L4 and L5 in rats^85^; we targeted recording to the L4 DRG because it bled less than the L5 DRG during surgery and recording. After the laminectomy, the rat was placed in a stereotaxic frame (Narishige) and its vertebrae clamped above and below the recording site to immobilize the spinal cord. Vertebral clamps were raised slightly to help mitigate movement of the DRG caused by breathing. The SCS lead was then implanted (see above). Rectal temperature was maintained at 37°C using a feedback-controlled heating pad (TCAT-2LV Animal Temperature Controller, Physitemp). Eyes were kept from drying by application of lube (Optixcare, CLC Medica). Hydration was maintained by intermittent i.p. injections of saline.

A multielectrode array (MEA) with 16 recording sites (A4 type, NeuroNexus) was implanted in the DRG. By recording in the DRG rather than directly from the DCs, as in ref. 17, we sampled neurons unbiased by the depth or medio-lateral positioning of their axon in the DCs (see above). The signal was amplified and digitized at 40 kHz with an Omniplex Data Acquisition System (Plexon). The signal was subsequently high-pass filtered at 250 Hz and spikes were sorted using the Plexon OfflineSorter. Single unit spikes had waveforms that were distinct from the SCS artifact and occurred with a latency of ∼2 ms; however, single unit spikes were often superimposed on the ECAP originating from multiple units and, for SCS ≥500 Hz (i.e. interpulse intervals ≤2 ms), spikes were superimposed on stimulus artifacts, which complicated sorting. This was solved by subtracting the artifact and ECAP recorded on many contacts from the single-unit spikes recorded on a single contact (see Supplementary Fig. 2).

Using SCS pulses applied at 2 Hz, the MEA was slowly lowered into the DRG while monitoring for SCS-responsive units. Once a so-called 2Hz+ unit was identified, the MEA position was adjusted to optimize recording. Next, the cutaneous receptive field was searched for by brushing the hind limb. SCS was then applied at ≥50 Hz. Units identified during ≥50 Hz SCS that were unresponsive during the 2 Hz search phase were labeled 2Hz-units; their receptive field was searched for during epochs of SCS not intended for other analysis. Most animals were tested at a single SCS frequency (50, 500, 1000, or 5000 Hz) because of the long (2 hour) duration of the standard protocol, which was designed to test whether neurons continued to respond to SCS. To compare responses in the same neuron to different SCS frequencies, a shortened protocol was applied in some rats.

#### Immunohistochemistry

In a separate set of rats, a single SCS protocol (c-SCS, kf-SCS, or sham) was applied for two hours with pulse amplitude set to 0.5 mA, which is ∼50% MT in most rats (see Supplementary Fig. 1). MT was not tested in these rats to avoid activating pERK. Five rats were prepared for each SCS protocol but one rat from the kf-SCS group and one in the sham group were excluded because of poor fixation. No electrophysiological recordings were conducted; instead, rats were perfused intracardially with fixative (4% paraformaldehyde in 0.1 M phosphate buffer (PB), pH 7.4) immediately after stopping SCS. L4 and L5 DRG were removed and stored overnight in fixative (same as above), and were then transferred to 30% sucrose in 0.1 M PB solution and stored at 4°C. To prepare sections, each DRG was embedded in an optimum cutting temperature medium (OCT; Tissue Tek), and 14‐μm sections were cut at –20°C on a cryostat (Leica) and collected on slides. Double‐labelling immunofluorescence was performed to observe expression of pERK in myelinated neurons identified by Neurofilament 200 (NF200) expression. DRG sections were initially permeabilized with 50% ethanol for 30 minutes, followed by a 10‐minute incubation in a 0.3% hydrogen peroxide solution. Sections were washed in PBS containing 0.2% Triton-X (PBS‐T) for 30 minutes between all incubations. Nonspecific binding of the secondary antibody was blocked by treating sections with a 10% normal goat serum (NGS) solution, diluted in PBS‐T, for 1 hour. Sections were then incubated overnight at 4°C with a rabbit polyclonal anti‐pERK antibody (Cell Signaling; 9101L, lot #30, 1:1000), and mouse anti-NF200 (Sigma; N0142, lot: 037M4779, 1:500) diluted in PBS‐T. After primary antibody incubation, sections were washed for 30 minutes in PBS‐T and incubated for 90 minutes in a biotin-conjugated goat anti‐rabbit IgG (Vector BA-1000; 1:400). After 30 minutes of PBS-T washes, sections were incubated with Vectastain Elite A+B reagents (1:250). Further signal amplification was achieved by treating the sections with tyramide (Perkin‐Elmer, Norwalk, CT; 1:75) for 7 minutes. Finally, sections were incubated in streptavidin-conjugated to Alexa Fluor 568 (Molecular Probes; S11226, 1:200) and goat anti mouse conjugated to Alexa Fluor 488 (Invitrogen; A11029, 1:800) for 2 hours. After incubation, sections were washed for 20 minutes in PBS and coverslipped with Aquapolymount (Polysciences, Warrington, PA). At least 5 sections per DRG were imaged using a Zeiss imager M2 microscope equipped with an AxioCam 512 mono camera and the ZEN Blue Imaging software (version 2.3; Zeiss). For representative images, image acquisition was performed using a LSM880 Confocal Microscope (Zeiss). Qualitative Image analysis was completed using ImageJ software by a blind experimenter. For this, DRG images were rank ordered, from DRGs containing the most pERK-expressing NF200-positive cells to those with the fewest pERK-expressing NF200-positive cells; those in the bottom third of the ranking were deemed pERK negative.

#### GCaMP-based calcium imaging

To express the genetically encoded calcium indicator GCaMP6f in DRG neurons, 5 µL of AAV9-CAG-GCaMP6f (Canadian Neurophotonics Platform Viral Vector Core Facility, Quebec, Canada) was injected intradermally in neonatal rats (<3 days old) as previously described^86^. Imaging was conducted 6-8 weeks later with two-photon microscopy using a VIVO 2-photon system (Intelligent Imaging Innovations, Denver, CO) comprising a AxioExaminer microscope (Zeiss), tunable Ti:Sapphire laser (Chameleon Ultra II, Coherent) set at 920 nm, and Slidebook 6 software. 2D full field scans were acquired at 6.2 Hz, 3D volume scans were acquired at 0.8 Hz. Each SCS protocol was applied continuously for 10 or 24 minutes. For the former, images were acquired continuously before, during and after SCS; for the latter, images were collected for several minutes before and after SCS onset, and again for several minutes before and after SCS offset. Anesthesia and surgical preparation were the same as for electrophysiological recordings. In a subset of experiments, the sciatic nerve was electrically stimulated (as for SEP recordings, see below) to gauge the number of cells activated for comparison with SCS-mediated activation.

#### Primary somatosensory (S1) cortex recordings

To record the somatosensory evoked potential (SEP) in rats, the SCS lead was inserted as described above but the remainder of the spinal surgery was minimized. Instead, the rat was placed in a stereotaxic frame with its head fixed by ear bars, a small cut was made to expose the skull and a hole was drilled 2.0 mm caudal to bregma and 2.7 lateral (right, i.e. contralateral to the sciatic nerve being stimulated) from midline. The underlying dura was removed. A four-electrode array (A4 type, NeuroNexus) was inserted to a depth of 200-500 μm. A wideband signal was recorded as described above for DRG recordings, but was subsequently low-pass filtered at 250 Hz. To evoke the SEP, the sciatic nerve was exposed and silver hook electrodes were used to apply electrical stimuli (monophasic pulses, 300 μs, 0.5 mA) at irregular intervals between 20 and 40 s using a constant-current isolated stimulator (model DS3, Digitimer). Stimulation was controlled by a second computer using a Micro1401 A-D board and Signal software (Cambridge Electronic Design) and was synced to the acquisition using the Omniplex system (Plexon). Ten nerve stimuli were applied before, during and after each SCS protocol. The order of SCS protocols was randomized across rats and each protocol was separated by 30 minute breaks during which the nerve was kept moist with saline.

#### H reflex measurements

To examine effects of SCS on the H reflex in rats, the medial plantar nerve was exposed over the left ankle. The nerve was stimulated as described above for the sciatic nerve for SEPs. The EMG was measured from plantar muscles with a pair of home-made steel needle electrodes. EMG responses were amplified (model DAM80, World Precision Instruments) and recorded through the same Micro1401 used to control stimulation. For medial plantar nerve stimulation, we used the lowest amplitude capable of triggering a maximal H wave and a stable M wave. Ten nerve stimuli were repeated before, during and after each SCS protocol. SCS protocols were tested in random order, like for SEP testing.

### Pig experiments (ECAP measurements)

All procedures in pigs were approved by Boston Scientific’s IACUC. Dorsal epidural spinal recordings were performed during concurrent SCS in six healthy pigs (*Sus scrofa domestica*, 40-65 kg) of both sexes in acute terminal experiments. Animals were initially sedated using propofol (≤6 mg/kg IV) and anesthesia was maintained using buprenorphine (1-10 µg/kg/hr IV). Once anesthetized, a 16-contact percutaneous lead (Infinion, Boston Scientific Neuromodulation [BSN]) was introduced via Tuohy Needle into the epidural spinal cord and positioned under fluoroscopic guidance at approximately midline underneath the lower thoracic vertebrae, between T10 and T14. Leads were then connected to a SCS External Trial Stimulator (BSN) that was, in turn, connected to a Tucker-Davis Technology (TDT) Physiological Recording System 3 (TDT, Alachua, FL) via a custom-made interface box (BSN). Biphasic symmetric SCS was delivered using a simple bipolar geometry (single anode and single cathode separated by two contacts, i.e., 8 mm) at 150 Hz or 1 kHz using a pulse width of 50 μs/phase for at least 10 minutes. Pulse amplitude was adjusted to the minimum level (within 0.1 mA) required to produce ECAPs visibly identifiable approximately 36 mm away from the cathode. Recordings were taken continuously from up to 32 channels using the TDT data acquisition system through all stimulation sessions. To enable peri-stimulus comparisons between 150 Hz and 1 kHz, the 1 kHz program was configured to stop for 6.67 ms every 500 ms. Recordings were sampled at 48,828 Hz, filtered using a 2^nd^ order Butterworth filter (high and low pass bands at 3 Hz and 8 kHz, respectively), and converted into MATLAB (R2019b, Mathworks, Natick, MA) readable format using a script provided by Tucker-Davis.

MATLAB files were segmented in two steps. First, locations of the first onset edges of all stimulation artifacts over the entire length of the experimental recording were identified in an arbitrary channel and reconciled with a record of amplitude change commands recorded by the Boston Scientific Clinician Programmer so that the recording could be synchronized with the initiation and cessation of stimulation. Continuous traces were converted into peri-stimulus intervals by time-aligning stimulation artifacts, collating peri-stimulus traces, and averaging when appropriate. ECAPs were identified based on characteristic morphology, namely a first negative (N1) and second positive (P2) peak with magnitudes at least 25% above the noise floor of the recording were detectable on multiple channels, and that N1 latency differences across multiple channels translated into physiological conduction velocities, as previously reported^62^.

### Human experiments (ECAP measurements)

All human experiments were approved by Western IRB. Representative ECAP recordings during 150 Hz and 1 kHz stimulation were taken from a single consented chronic pain patient who was concurrently undergoing a SCS trial. The test device system used to record ECAPs consisted of a passive custom-designed breakout box (BSN) that enabled connection between a commercially available SCS external trial stimulator and trial implanted percutaneous leads (BSN) to the TDT data acquisition system described above. SCS parameters, data acquisition and filtering were the same as described above except that pulse amplitude was adjusted to just above the threshold at which the subject could feel paresthesia during 150 Hz SCS. ECAPs were analyzed as described above.

### Computational modeling

All computer code will be made available at ModelDB upon publication.

#### Phenomenological model of SCS-evoked spiking

To test if time-dependent changes in excitability are sufficient to explain patterns of SCS-evoked spiking, we modeled spike generation as a statistical process. A random number is drawn from a Gaussian distribution (mean, *µ*; standard deviation, *σ* = 0.05) and produces a spike if it exceeds threshold. The random number is drawn at the interpulse interval, namely every 20 ms for c-SCS (=1/50 Hz) vs every 1 ms for kf-SCS (=1/1 kHz). Importantly, the Gaussian distribution and threshold evolve in a time-dependent manner to reflect fluctuations in post-spike excitability.

Fluctuations in excitability were modeled as a fast refractory period (referred to here as an afterhyperpolarization, or AHP, although sodium channel inactivation rather than hyperpolarization may be responsible), an intermediate afterdepolarization (ADP), and slow adaptation. The AHP and ADP are each modeled as 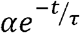, using different values for scaling factor α and time constant *τ*, and are summed. These two processes, which are relatively fast and are reset after each spike, shift the Gaussian distribution from which random numbers are drawn. To account for slow adaptation, threshold is incremented by Δ*Th* each time a spike occurs and then slowly relaxes according to *τ*_adaptation_. The decay is slow enough that changes in threshold accumulate across spikes, which can lead to bursting (see below). The fast and slow changes are modeled separately to help depict their interaction.

At the onset of SCS, threshold is defined as 0 and *µ* = 0. Random numbers are drawn (at the interpulse interval) until one exceeds 0, thus producing the first spike. When a spike occurs, the Gaussian distribution is shifted downward (by the AHP), which reduces the probability of random numbers exceeding threshold shortly after the preceding spike. Over time, the Gaussian distribution shifts upwards (as the AHP gives way to the ADP), thus increasing the probability of random numbers exceeding threshold. The ADP makes it likely that another spike will occur between ∼3 and 10 ms after a preceding spike. However, each time a spike occurs, the threshold is incremented. Accumulation of those increments across several spikes (i.e adaptation) competes with the increased likelihood of spiking during the ADP, eventually terminating the burst of spikes when a spike fails to occur during an ADP. Effects of the fast processes (i.e. the AHP and ADP) are only evident when the model is stimulated at short intervals (as during kf-SCS); stimulating at longer intervals (as during c-SCS) fails to reveal the rich dynamics since the system has effectively returned to baseline before the next pulse. All excitability parameters (*α*_AHP_, *τ*_AHP_, *α*_ADP_, *τ*_ADP_, Δ*Th*, and *τ*_adaptation_) were varied randomly across models to introduce cellular heterogeneity. Values and additional methodological details are available in the code.

#### Biophysical model of human DC axons

A single 14.0 μm diameter myelinated fiber consisting of 10 nodes of Ranvier and a point source were placed in a homogeneous, isotropic environment (σ = 0.2 S/m). The point source was positioned 1 mm to the side of the first node. Axon biophysics and geometry were largely retained from a previously described double-cable model of a myelinated peripheral axon^87^, but two modifications were made. First, a slow afterhyperpolarizing potassium conductance (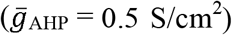), with a state variable *z* and time constant τ = 200 ms, was implemented within the nodes consistent with Hodgkin-Huxley formalism, as previously described^88^:

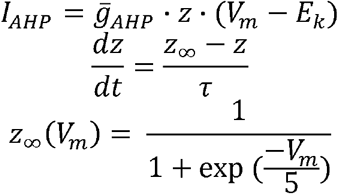

Second, the time constant of the fast sodium channel state variable *h* was slowed down by a factor of 2.25 to account for the possibility of fiber adaptation to high frequencies of stimulation.

In addition, Ornstein-Uhlenbeck noise (mean = 0, SD = 8 pA, τ = 0.5 ms) was injected into each node to mimic channel noise. All nodes on a given axon received the same noise. Concurrently, a pulse train *I*_0_ consisting of symmetric, biphasic rectangular pulses (same waveform as for experiments; see Fig 1A) was delivered at 50 Hz or 1 kHz through the point source. Changes in extracellular voltage *V*_e_ thus produced were applied to all *n* segments of the model axon over each time point *t*, as

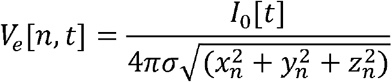

Where σ represents isotropic bulk extracellular conductivity^89^ . Effects of the extracellular waveform were coupled to the axon membrane using the “extracellular” mechanism in NEURON. All simulations were conducted using NEURON (v7.7.2) and using the Crank-Nicolson integration method with a time step of 0.005 ms.

Simulations were conducted in 30 models in each SCS condition. Each model received different noise. Sodium conductance was varied between 2.0 to 3.0 S/cm^2^ in 0.25 S/cm^2^ increments to account for cellular heterogeneity and stimulus amplitude was varied between 0.16 and 0.24 mA in 0.02 mA increments to account for variable stimulation (e.g., distance from the electrode, tissue conductance). Responses were aligned to mimic a set of 30 simultaneously tested axon models.

#### Model of human spinal cord and ECAP simulations

A previously published three-dimensional finite element model (FEM) of a human thoracic spinal cord built using COMSOL Multiphysics (Version 5.1, COMSOL Inc., Burlington, MA) was used to calculate electrical tissue responses to 50 Hz and 1 kHz SCS^90, 91^ using simple bipolar (single anode, single cathode, separated by 8 mm) stimulation from a percutaneous lead placed at the midline of the cord. The geometry of the FEM was generated by extruding a coronal spinal cord cross-section that was, itself, traced from an image of a cadaveric cross-section of the low thoracic spinal cord (T8-T9)^92^. Tissue and SCS electrode properties (dimensions, lead positions, conductivities) were preserved but the total rostrocaudal length of the FEM was extended to 200 mm to accommodate dorsal column fibers long enough that action potentials take >2 ms to propagate from the activation site (under the stimulation cathode) to the boundary of the model. Continuous boundary conditions were used to define internal boundaries of the model beyond those specified. Dirichlet boundary conditions (V = 0) were used to define the outer boundaries of the model, and extracellular voltages at steady state, assuming a quasi-static tissue medium^93^, resulting from stimulation were computed by solving for Laplace’s equation (∇·_σ_∇V = 0).

Extracellular voltages calculated from the finite element model were coupled to biophysical cable axons (see above) seeded in the spatial model uniformly on a 200 µm (rostrocaudal) × 100 µm (dorsoventral) grid down to 1 mm in depth from the dorsal surface of the spinal white matter to enable simulation, using NEURON, of dorsal column axon responses to 50 Hz and 1 kHz SCS. Axon diameters ranged from 5.0 µm to 14.0 µm in 0.5 µm increments; geometric properties of fiber diameters not originally included in McIntyre et al. were derived in MATLAB using piecewise cubic Hermitian interpolation. Axons were modeled as described above but with the noise applied at each node scaled by fiber diameter so that noisy voltage fluctuations were roughly equivalent across axons of different diameters.

Membrane current responses at each node were then converted to extracellular potentials as would be recorded from a percutaneous SCS lead contact through transfer impedance matrices for each SCS contact derived from the spatial model, based on Helmholtz Reciprocity Theorem and consistent with previously described methods^64^. Following such conversion, individual axon responses were scaled by expected fiber densities in human spinal cord by diameter^94^, time-aligned to stimulation, and summed to form a simulation of the extracellular dorsal column ECAP on a given contact. For ECAP simulations, axons were not injected with Ornstein-Uhlenbeck noise due to deleterious effects of noise on the extracellular signal.

#### Model of feedforward inhibitory (FFI) circuit

The circuit included 10 cells each of three cell types – input neurons (primary afferents), inhibitory neurons, and output neurons – connected all-to-all (see below). Input neurons were modeled simply as experimentally recorded spike trains evoked by c-SCS or kf-SCS (see Fig. 2A) and selected to have a group average firing rate of ∼50 spk/s. Inhibitory neurons and output neurons were modeled as leaky integrate-and-fire neurons. Both of these cell types had similar parameters (*C*_M_ = 50 pF, τ_ref_ = 0.1 ms, τ_syn_ = 3 ms, *V*_reset_ = -85 mV, *V*_thresh_ = -51 mV), but differed in their membrane time constants (τ_mem_) based on previous models of a thalamocortical FFI circuit^95^. Specifically, to introduce heterogeneity into these models, each cell had a slightly different τ_mem_ drawn from a Gaussian distribution centered at 10 ms for inhibitory cells and 16 ms for output cells; standard deviation was 1 ms in each case. All inhibitory cells and output cells also received independent Poisson background noise using independent NetStims. All inhibitory neurons and output neurons received an EPSP each time an input neuron fired, and each output neuron received an IPSP each time an inhibitory neuron fired (with a 2 ms delay). Excitatory synaptic weights were 120 and inhibitory weights were varied between 0 and -200. Simulations were performed in NetPyNE^96^.

## ACKNOWLEDGEMENTS

This work was funded by a grant from Boston Scientific Neuromodulation, a Foundation Grant from the Canadian Institutes of Health Research, and a John Edwards Leadership Fund award from the Canadian Foundation for Innovation to SAP. We thank Russell Smith and Manon St-Louis for expert technical assistance.

## CONTRIBUTIONS

The project was conceived by SR and SAP. Electrophysiology was carried out by BS, immunohistochemistry by NY, calcium imaging by BS and SR, and ECAP measurements by TZ. Computational simulations were carried out by NA, TZ, LM with input from DA-B. DA-B also assisted with data analysis. SAP, SR, AR-d-S, and RE supervised all research. SA prepared the final manuscript with input from all authors.

## SUPPLEMENTARY FIGURES

**Supplementary Figure 1.**
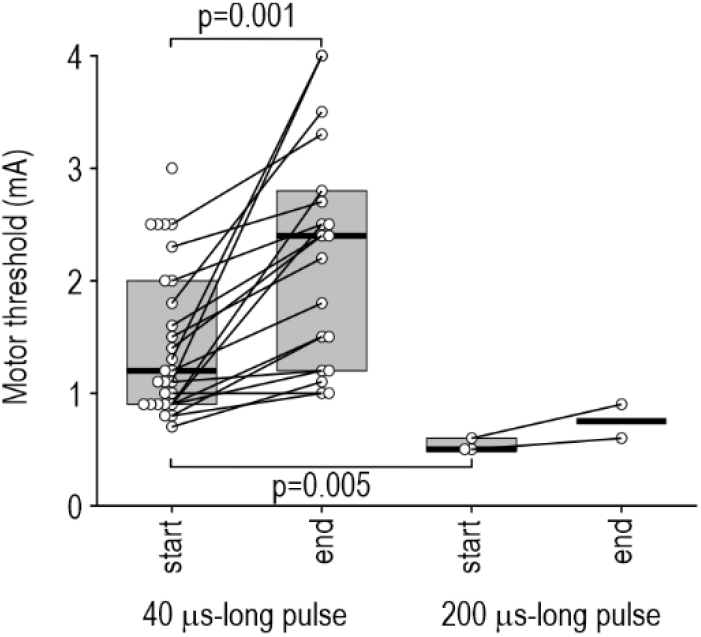
Motor threshold (MT) in rats. MT was determined during 2 Hz SCS. 40 µs-long pulses (see Fig. 1A) were used for all SCS frequencies except in a subset of experiments where c-SCS was re-tested using 200 µs-long pulses. Graph shows MT values from all rats. Boxes summarize median and interquartile range for each condition. For 40 µs-long pulses, median MT increased from 1.1 mA (0.9 – 1.575; interquartile range) to 2.4 mA (1.275 – 2.775) between the start and end of experiments (*Z*=3.72, *p*<0.001, Wilcoxon signed rank test). This likely explains why some units stopped spiking during sustained SCS, and why spiking could be restored by a small increment in pulse amplitude (see Supplementary Fig. 4). The starting MT for 200 µs-long pulses (0.5 mA, 0.5-0.575) was significantly less than for 40 µs-long pulses (*U*=0, *p*=0.005, Mann-Whitney test). Notably, the smaller amplitude (∼0.5×) did not fully offset the longer pulse width (5×), meaning substantially more charge (current × time) was applied with 200 µs-long pulses than with 40 µs-long pulses. Supplementary Discussion – Stimulus conditions must be considered when comparing MT values. For example, Song et al.^15^ measured MT using monophasic, 200 µs-long pulses delivered at 4 Hz. Testing with low frequency, 200 µs-long pulses yielded MT values (0.6-0.8 mA) similar to us (0.50-0.58 mA). But Song et al. tested high-frequency SCS (0.5, 1, and 10 kHz) using 24 µs-long pulses; MT is much higher for a pulse width nearly an order of magnitude shorter, so saying that short pulses of 0.3-0.4 mA were 40-50% of MT is misleading and may explain why Song et al. found no electrophysiological consequences of high frequency SCS. Shechter et al.^28^ measured MT using biphasic 24 µs-long pulses delivered at 4 Hz, and used the same pulse width for all SCS frequencies, like we did; they also observed an onset response for high frequency SCS at ≥80% MT. Measuring the compound motor action potential in humans during electrical nerve stimulation, Deletis et al.^97^ found that threshold was reduced by 6-42% during trains of 0.5-10 kHz, respectively, compared with single pulses. Dietz et al.^98^, testing awake animals with a pulse width similar to Shechter et al., reported MT values much smaller than Shechter et al. (0.27 vs ∼0.8 mA). Such differences boil down to technical factors some of which, like lead placement^99^, are impossible to fully standardize. Expressing SCS intensity relative to MT helps ensure application of a similar effective “dose” despite those differences. Anesthesia is another important consideration. Dietz et al.^98^ recently showed that isoflurane (1.5-2.5%) increased MT by ∼5.5× compared with awake rats (without causing a concordant increase in ECAP threshold). Isoflurane (1.5%) was previously reported to double MT^100^. In contrast, urethane has only modest effects on subcortical processing, including reflexes^101^; for instance, Angel and Unwin^102^ reported that “cells in the cuneate nucleus…showed no reduction in their probability or latency of discharge to supramaximal stimuli, until the anaesthetic depth [of urethane] seriously impaired the animal’s respiratory effort.” Cortical processing of somatosensory input is affected by urethane^103^ but less so than by isoflurane^104, 105^. In short, urethane is unlikely to increase MT and, more importantly, does not cause a discordant increase in MT and ECAP threshold (see below). Other experimental factors must also be considered when considering the absolute value of MT; for example, minimizing breathing movements for stable single unit recordings by lifting the vertebrae (see Methods) enlarges the epidural space, decreasing current delivery to the cord^80^ and thus increasing MT, in the same way posture affects SCS in patients. Normalizing against MT is also useful for gauging other thresholds. Shechter et al. found that sensory threshold (ST) was ∼50% of MT in awake rats. Dietz et al. found that ECAP threshold was ∼33% of MT in awake rats. These relative values align reasonably well even if the absolute values (∼0.4 mA and 0.09 mA, respectively) do not; the discrepancy is mostly accounted for by the difference in MT (see above). We similarly started to detect ECAPs at 30-40% MT (see Fig. 5A). Notwithstanding some discrepancies (e.g. ref. 100 detected ECAPs at 71-87% of MT), there is general agreement that ECAP and sensory thresholds sit somewhere between 30 and 50% of MT. For comparison, simulations in human spinal cord predict that the ratio of ST to discomfort threshold is anywhere from 23% to 63% based on electrode configuration^106^.

**Supplementary Figure 2.**
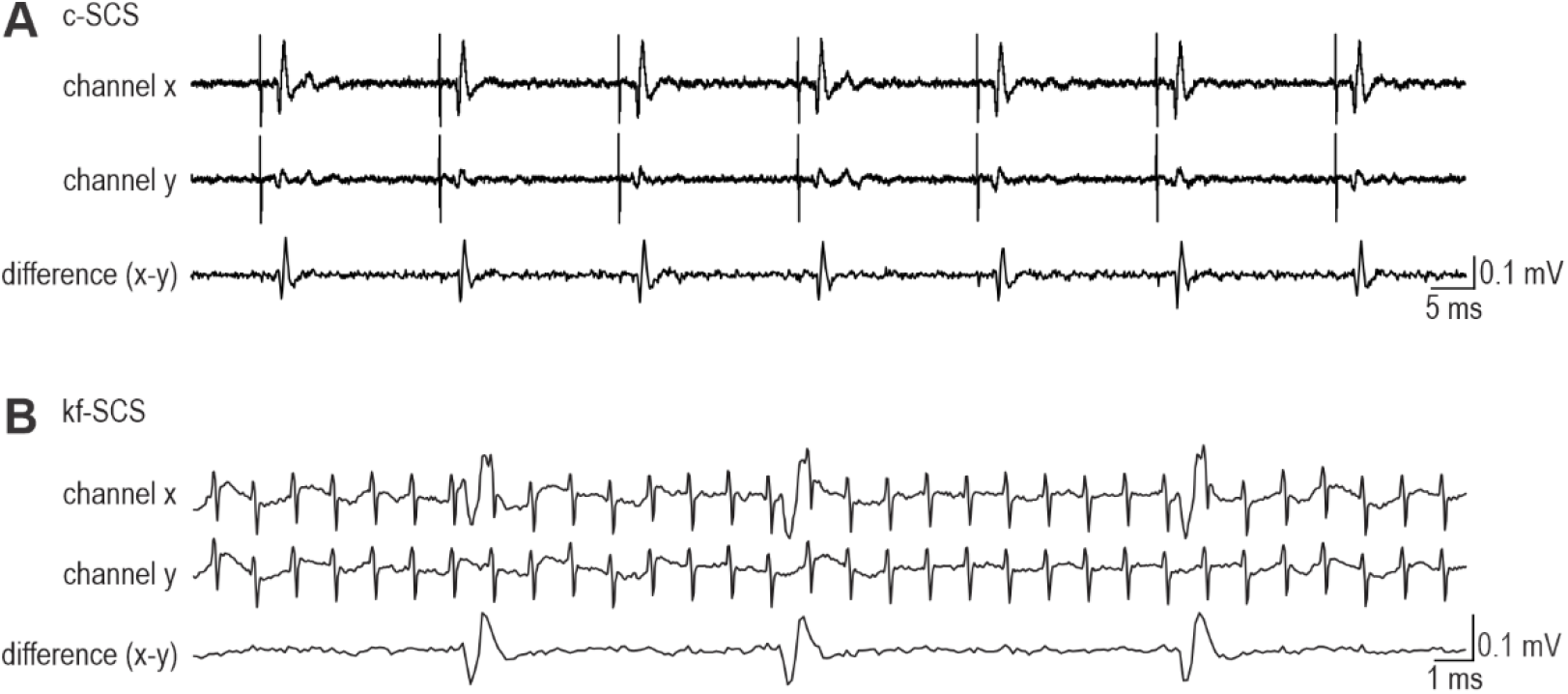
Isolation of single-unit activity. Multielectrode arrays (MEAs) were implanted into L4 DRG to record spikes conducted antidromically from the DC stimulation site (see Fig. 1A). Whereas stimulus artifacts and ECAPs were recorded on multiple contact, single-unit activity was detected exclusively by a single contact (presumably the one closest to the activated unit). Subtracting signals recorded from different electrodes (with similarly sized artifacts and ECAPs) yielded clean single-unit recordings, from which spikes were then sorted. **(A)** During c-SCS, ECAPs and single-unit spikes occur between artifacts, but the subtraction step nonetheless helps separate single-units spikes from ECAPs since they invariable overlap during c-SCS. See also Figure 5A. **(B)** During kf-SCS, single-unit spikes occur amidst artifacts and are difficult to sort without the subtraction step. On the other hand, ECAPs become small and irregular during kf-SCS and do not, therefore, obscure single-unit spikes.

**Supplementary Figure 3.**
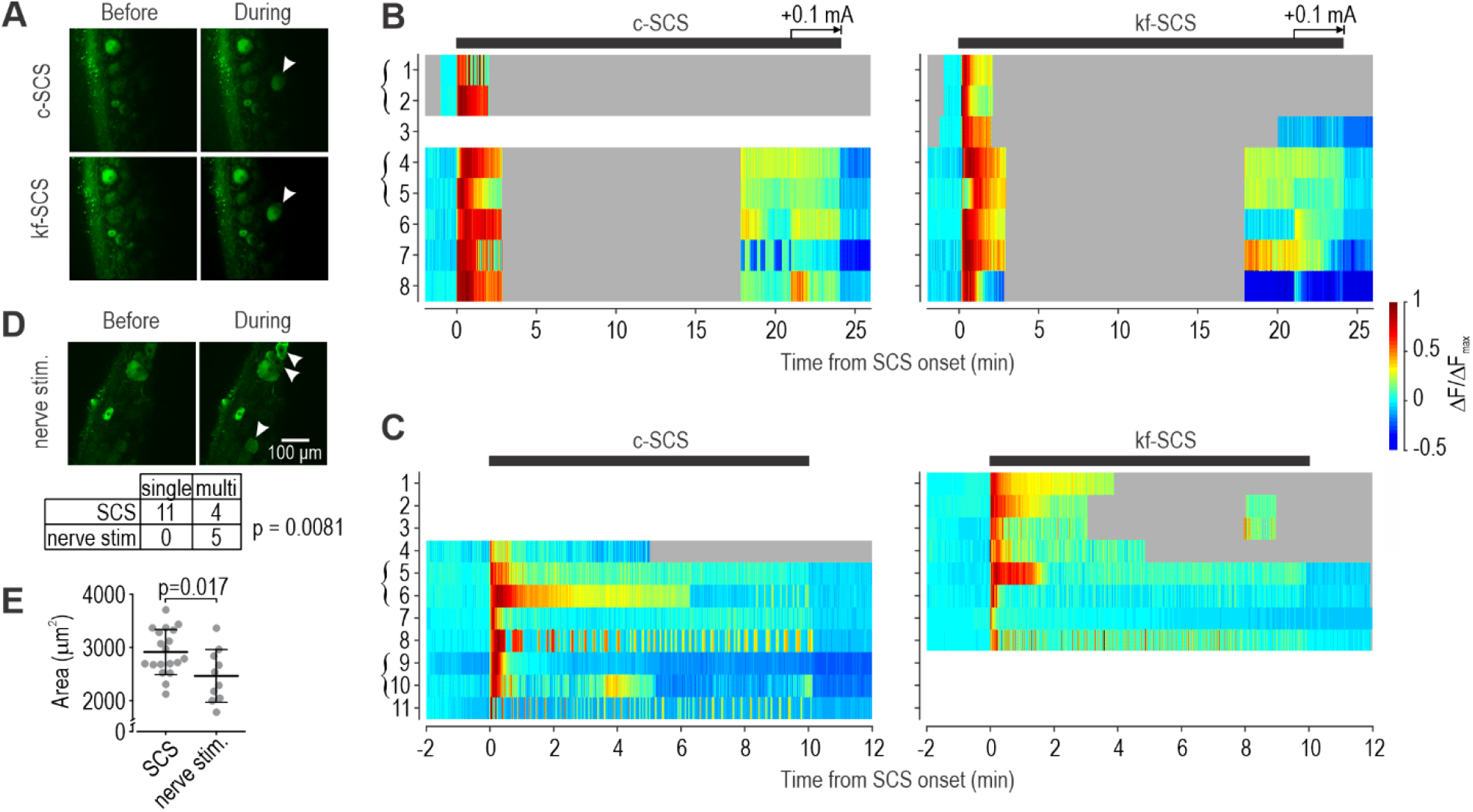
Additional GCaMP-based calcium imaging in rat DRG. All SCS was applied at 50% MT. Both forms of SCS were tested in most experiments, in random order. Normalized changes in GCaMP fluorescence (∆F/∆F_max_ = ∆F/F normalized by the maximal ∆F/F in that neuron) are plotted in color; gray and white indicate when imaging was paused or not conducted, respectively. Brackets indicate cells simultaneously imaged in the same field of view. **(A)** Sample images show activation of a single cell (arrowhead) during c-SCS (top) and kf-SCS (bottom). Non-responsive cells are also evident. **(B)** Eight cells were imaged at multiple z-positions at 0.84 frames/s from 3 rats during c-SCS (left) and kf-SCS (right) applied for 24 min. Imaging was conducted at the onset and offset of SCS. In all cells, ∆F/∆F_max_ increased at SCS onset and decreased at offset; incrementing SCS intensity shortly before its offset tended to increase fluorescence. For kf-SCS, pulse amplitude was ramped up at the onset to minimize motion artifacts caused by muscle contractions that were imperceptible to the naked eye. **(C)** An additional 11 cells were imaged at a single z-position at 6.25 frames/s from 4 additional rats. Many cells exhibited ∆F/∆F_max_ fluctuations consistent with bursting. **(D)** Sample field showing activation of 3 cells during electrical stimulation of the sciatic nerve (10 pulses at 10 Hz, 0.5-4 mA). Table summarizes number of fields imaged in which one or more cells were activated. Multi-cell activation was significantly more common during peripheral nerve stimulation than during SCS (*p*=0.0081, Fischer exact test), consistent with sparse activation during SCS. **(E)** Cross-sectional area of activated somata. Cells activated by SCS (2910±425 µm^2^; mean±SD) were significantly larger than those activated by nerve stimulation (2461±496 µm^2^) (*T*_27_=2.55, *p*=0.017, unpaired T-test). With a somatic diameter of ∼60 µm, SCS-activated neurons are amongst the largest in the rat DRG^107^, consistent with SCS selectively activating large DRG neurons (which are the only afferents that send axons up the DCs).

**Supplementary Figure 4.**
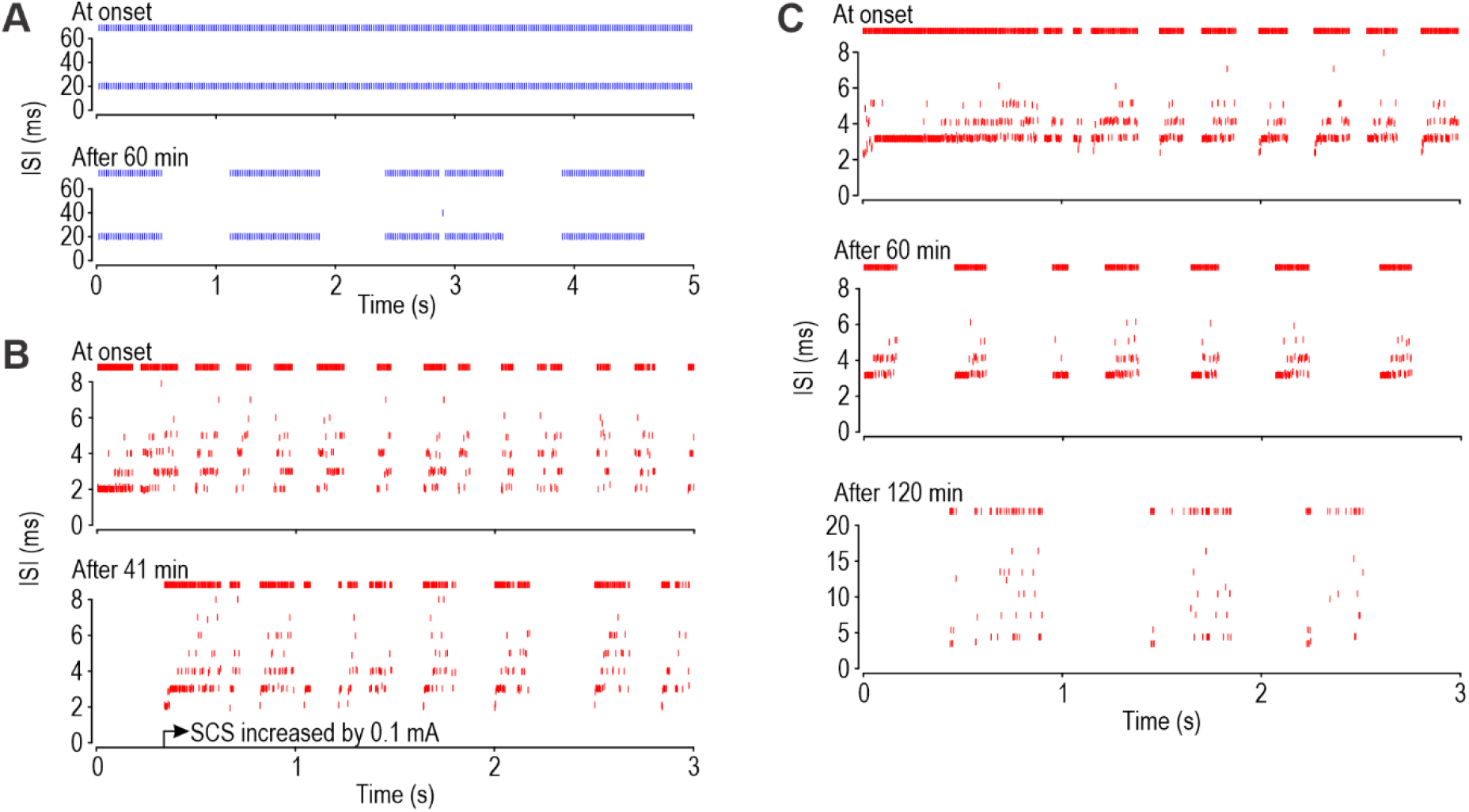
Differentially synchronized spiking persists during sustained SCS. Each ISI-vs-time plot shows data from a single unit, unlike in Figure 3 where data from many units are superimposed. Rasters are shown at the top of each panel. **(A)** Sample response during c-SCS. Tonic spiking at the onset of c-SCS (top) eventually gave way to bursting after one hour (bottom) but all intraburst spikes continued at the 20 ms interpulse interval. **(B)** Sample response during kf-SCS. Short bursts at the onset of kf-SCS stopped by 41 minutes of stimulation, but the unit resumed firing with a similar pattern when SCS intensity was incremented from 0.4 to 0.5 mA (bottom). In this rat, MT increased from 1.1 mA at the start of the experiment to 4.0 mA at the end. **(C)** Another sample response during kf-SCS. Rasters are shown at the top of each panel. Interburst intervals were initially quite short (top) but became longer after an hour of stimulation (middle). After another hour of stimulation (bottom), intraburst spikes occurred at a lower rate (note the change in y-axis scale) but still at multiples of the 1 ms interpulse interval.

**Supplementary Figure 5.**
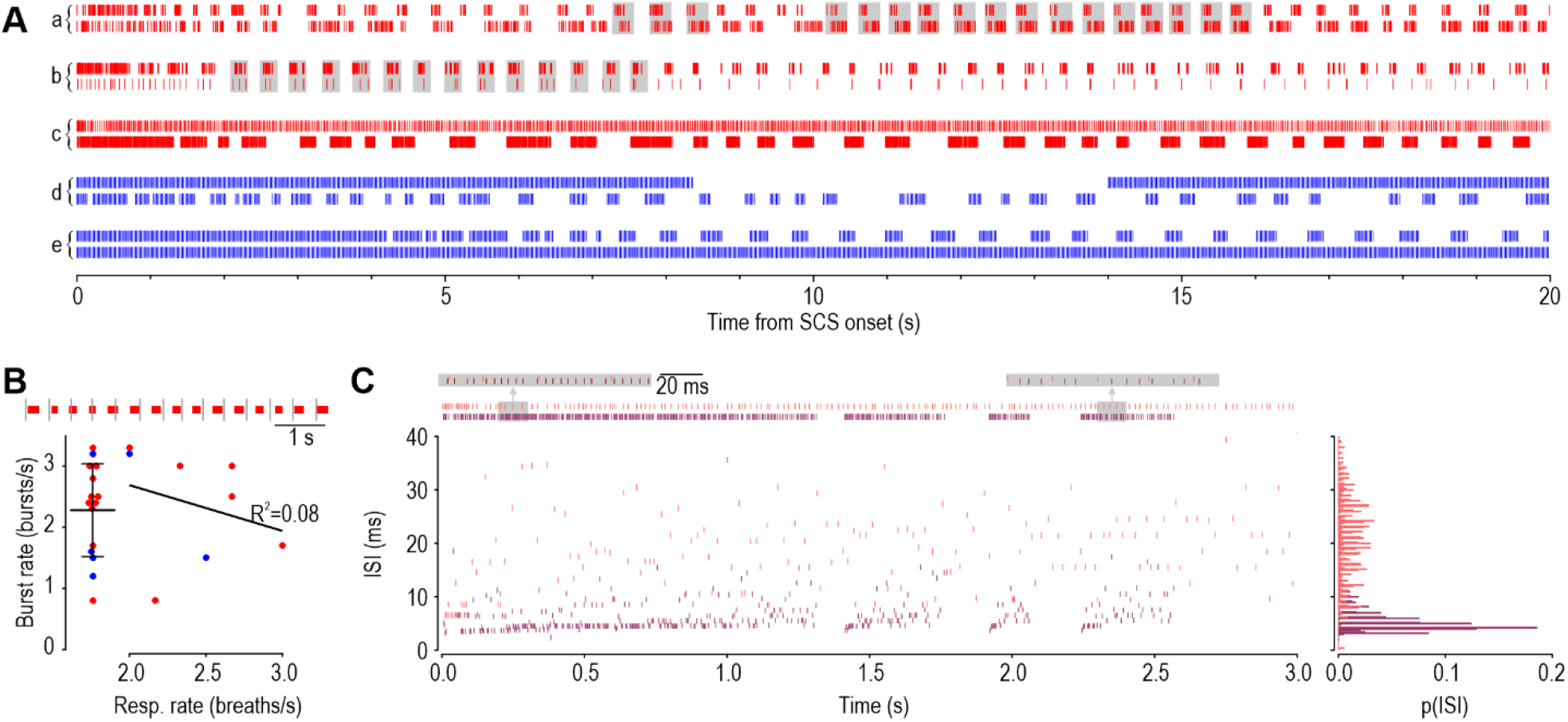
Intraburst spikes remain asynchronous even during coincident bursts. **(A)** Sample rasters from pairs of simultaneously recorded units during kf-SCS (red) and c-SCS (blue) showing that bursts often coincide (gray shading) but are imprecisely coordinated insofar as bursts in each unit start and stop at slightly different times, and comprise different numbers of spikes. Pair *a* (analyzed in Figure 3E,F) and pair *b* are similar in this regard, but differ from pair *c* (analyzed in panel C), where one unit bursts while the other spikes continuously, as commonly observed during c-SCS (pairs *d* and *e*). **(B)** Bursts occurred at a rate of 2.28±0.76 (mean±SD), which did not differ between c-SCS and kf-SCS (*t*_13_=1.27, *p*=0.23, unpaired t-test). Burst rate is similar to breathing rate (2-3 breaths/s) but they did not correlate (*R*^2^=0.08, *p*=0.49) in rats in which breathing was measured at the start of experiments. Raster at top shows that timing of breathes (grey bars) relative to burst onset slowly shifted (precessed) over time. This analysis was conducted in only a few rats, but in no case was a clear temporal relationship discerned, arguing that burst rate is intrinsic to each neuron. **(C)** Intraburst spikes never synchronized during kf-SCS. Data here are from a second pair of simultaneously recorded units analyzed like in Figure 3F. One unit fired continuously at a relatively low rate (pink) whereas the other fired in bursts (crimson). Both units fired at multiples of the interpulse interval but did so on different pulses (i.e. asynchronously).

**Supplementary Figure 6.**
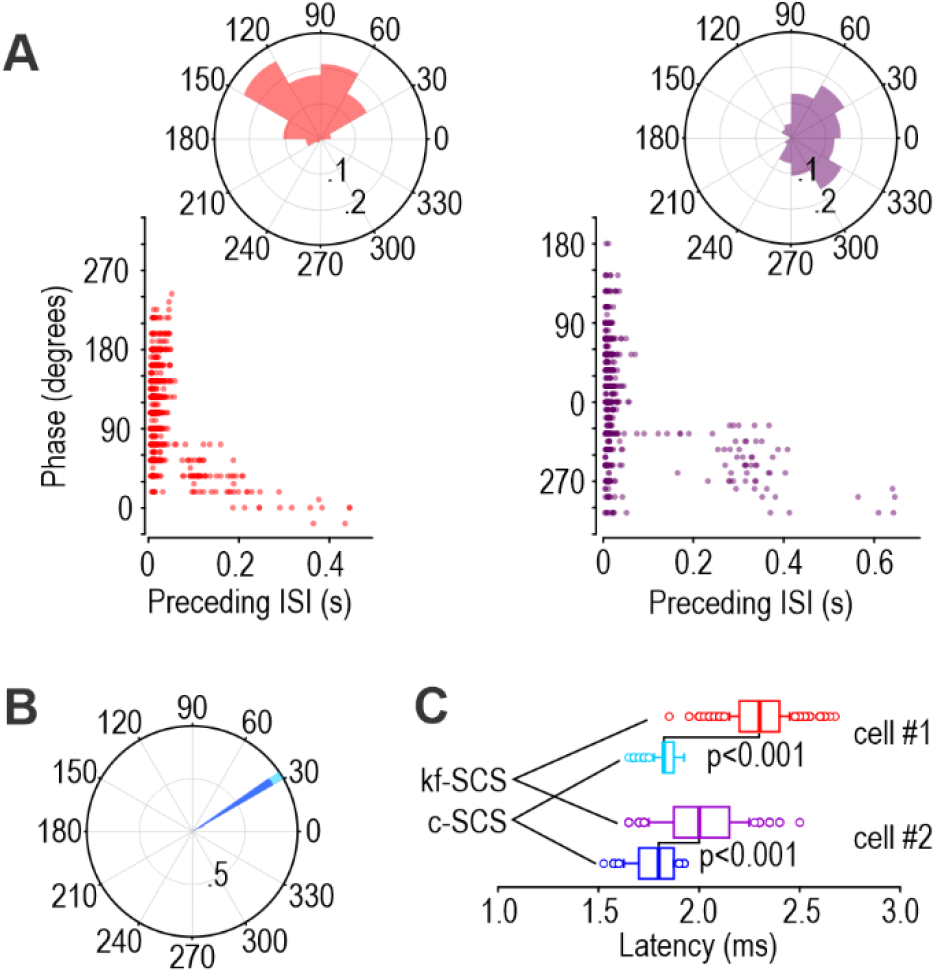
Temporal dispersion contributes little to desynchronization. **(A)** During kf-SCS, two simultaneously recorded units fired at slightly different phases relative to SCS pulses, according to polar plots (top). This means that if both units fired on the same pulse, the spike in the purple unit would typically occur 0.3 ms before the spike in the red unit (after converting phase to latency). Bearing in mind that spikes are recorded at a different location from where they originate, the difference in latency (*U*=60540.5, *p*<0.001, Mann-Whitney test) is not explained by the small difference in conduction velocity (16.7 vs 16.8 m/s based on 2 Hz testing). Moreover, any particular pair of spikes could exhibit an even larger difference given the broad distribution (high variability) of phases in each unit. Some of this variability is explained by previous spiking, as revealed by plotting phase against the preceding ISI (bottom). The first spike in a burst (i.e. preceded by a long ISI) had a shorter-than-average latency, suggesting that latency increases – most likely because conduction velocity slows^108^ – as the axon fatigues during a burst. **(B)** Polar plots for spikes from the same two units during c-SCS. The units now appear to spike at the same phase, but the longer stimulus period (20 ms during c-SCS vs 1 ms during kf-SCS) obscures differences in the absolute timing; in other words, latencies normalized by a long period (to calculate phase) appear less variable than if normalized by a short period. **(C)** Direct comparison of latencies shows, for both units, that latency was longer during kf-SCS than during c-SCS (U>700, p<0.001, Mann-Whitney tests) and more variable (i.e. spike time jitter is higher). But compared with the desynchronizing effect of firing on different pulses, which entails shifting the spike time by 1 ms for each skipped pulse (for 1 kHz SCS), jittering the spikes by fractions of a millisecond has a modest effect. That said, for stimulation frequencies ≥5 kHz, where interpulse intervals are ≤0.2 ms, jittering starts obscuring which pulse evoked the spike (see Supplementary Fig. 7C). The phase locking described above is equivalent to what Crosby et al.^17^ refer to as phase synchrony and measure with vector strength. When they report that *phase* synchrony is high during 1 kHz SCS, they mean that phase locking is strong, not that that unit fires synchronously with other units. Our results are entirely consistent with their vector strength analysis, but our definition of synchrony emphasizes whether units fire on the same SCS pulse, rather than whether they fire at the same phase (of potentially different pulses).

**Supplementary Figure 7.**
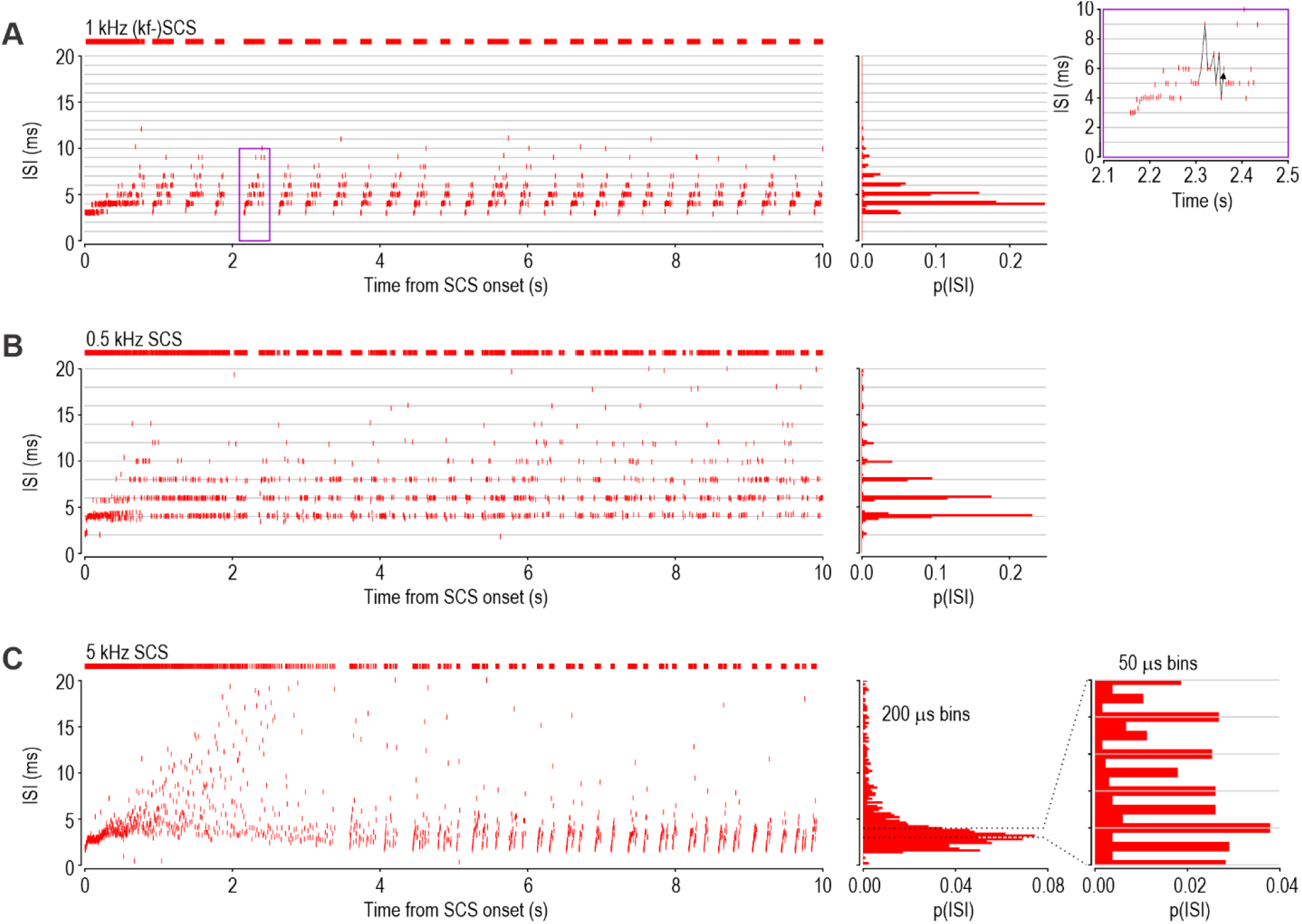
SCS at 0.5 kHz and 5 kHz evokes asynchronous spiking. Each panel shows data from a different 2Hz+ unit. **(A)** Sample response during 1 kHz SCS. Most spikes occurred with intervals between 3 and 7 ms, at integer multiples of the 1 ms interpulse interval (gray lines). Inset shows enlarged view during a typical burst (purple box) to highlight the evolution of intraburst ISIs: ISIs get longer over the course of each burst but the progression is erratic (see black arrow), consistent with stochastic spike initiation. **(B)** Sample response during 0.5 kHz SCS. Here, ISIs occur at multiples of the 2 ms interpulse interval. **(C)** Sample response during 5 kHz SCS. Most ISIs fell between 2 and 5 ms with an overall distribution (envelope) similar to that seen in panels A and B. Using our standard 0.2 ms-wide bin to construct the histogram was insufficient to resolve the individual peaks expected at multiples of the 0.2 ms interpulse interval. Inset shows an enlarged view of the histogram re-calculated using 50 µs-wide bins. ISIs occurred at integer multiples of the interpulse interval, but also in antiphase (i.e. halfway through the interpulse interval) for reasons that remain unclear. The key observation is that repetitive stimulation at intervals shorter than the axon refractory period caused axons to fire intermittently, on only a subset of pulses; indeed, the envelope for ISI histograms was similar across 0.5 to 5 kHz SCS. All evidence argues that pulses are skipped irregularly at these SCS frequencies.

**Supplementary Figure 8.**
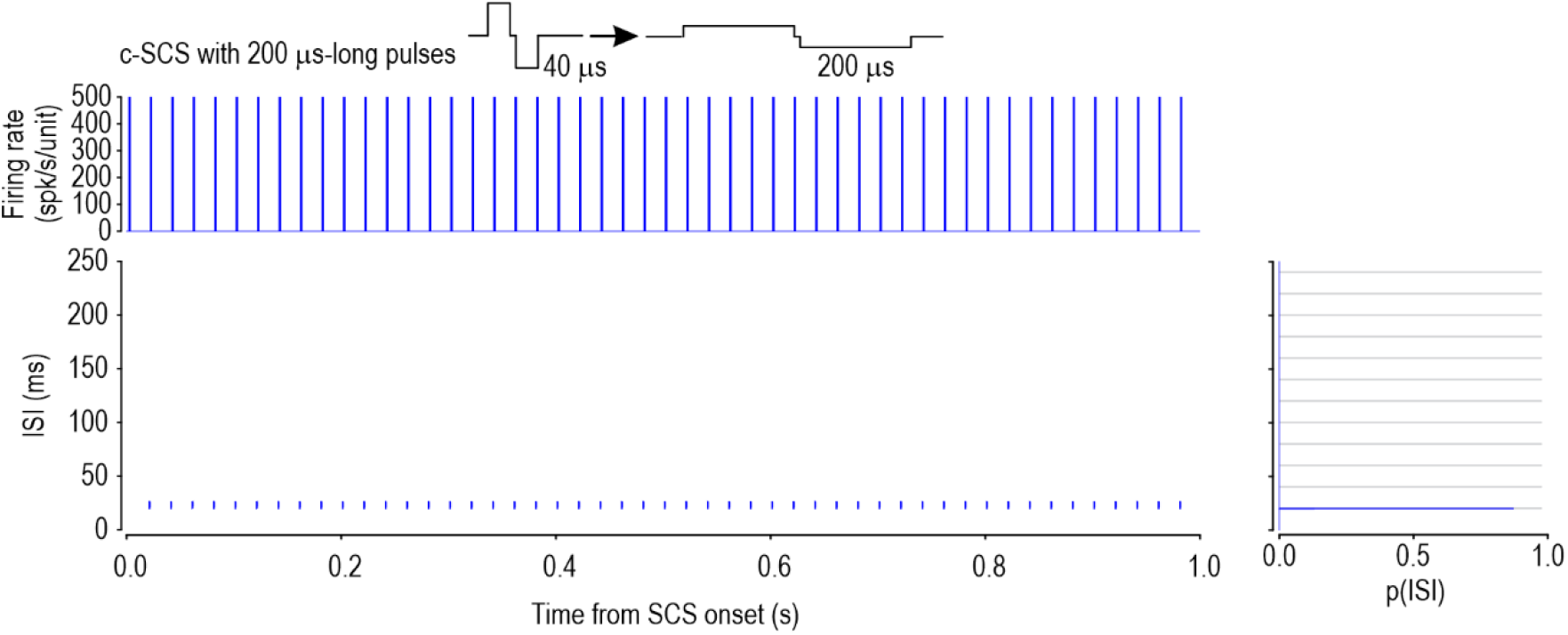
C-SCS using longer, lower-amplitude pulses still evokes synchronous spiking. Three units were tested with c-SCS (50 Hz) with pulse width increased to 200 µs (from 40 µs). Data from all units are included. As expected, MT was lower when using the longer pulse width (see Supplementary Fig 1). Conventional-SCS was tested at 50% of this lower MT. Data, plotted like in Figure 3, show that virtually all spikes occurred at the 20 ms interpulse interval, which translates into synchronized spiking.

**Supplementary Figure 9.**
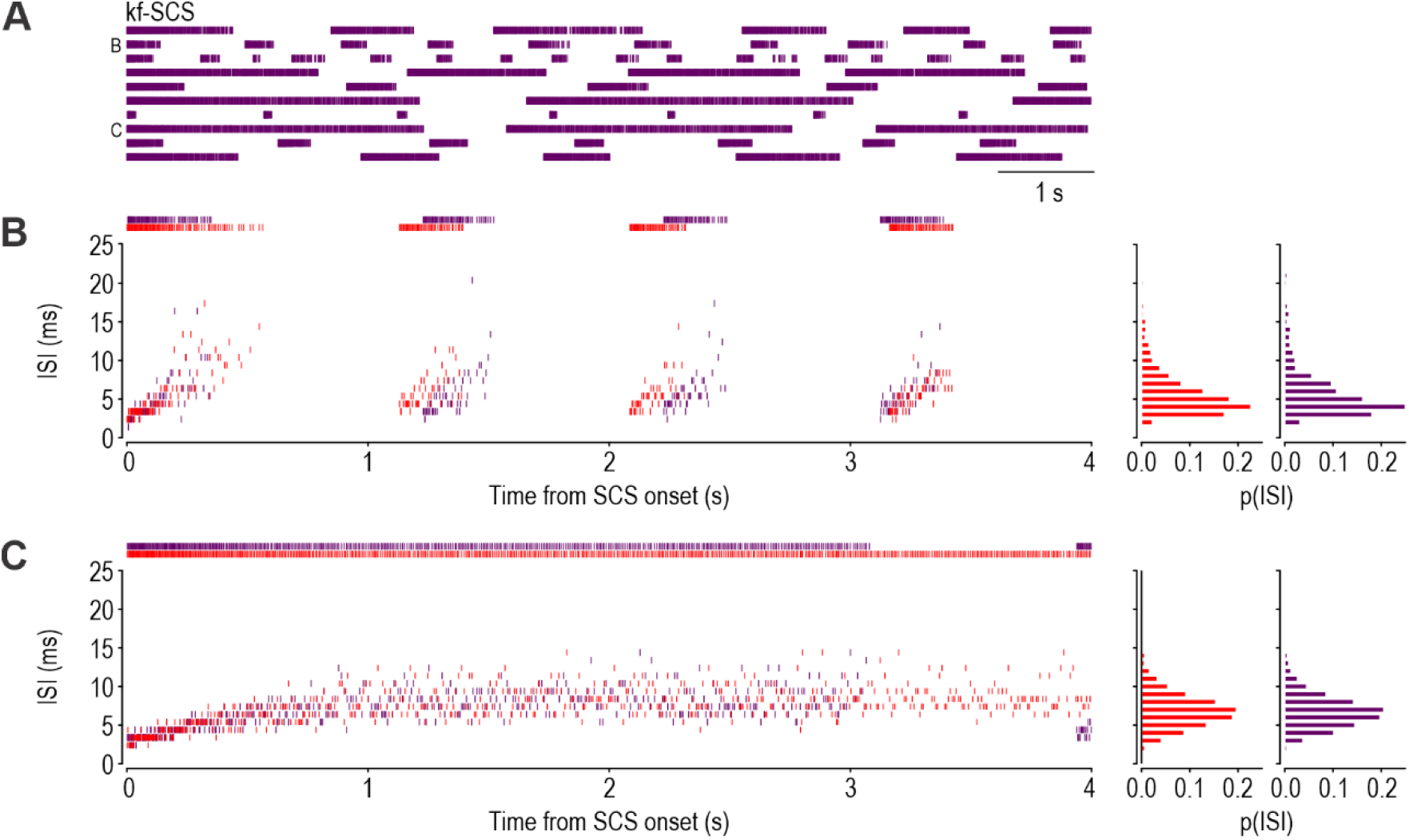
Effects of uncorrelated noise on spiking in phenomenological model. **(A)** Using the same 10 model axons as in Figure 4B, kf-SCS simulations were re-run with different noise. Rasters marked B and C are compared (in panels B and C, respectively) with corresponding rasters in Figure 4B, i.e. for different noise in an identical axon. **(B)** The same model axon given different noise produced similar ISI distributions and bursts, but the timing of individual spikes and bursts differed. **(C)** Equivalent analysis in a different axon model yielded comparable results. Comparing with experimental data (see Fig. 2A), these results argue that noise affects the timing of bursts and intraburst spikes, but that differences in intrinsic axon properties account for variations in burst properties (e.g. number of spikes per burst).

**Supplementary Figure 10.**
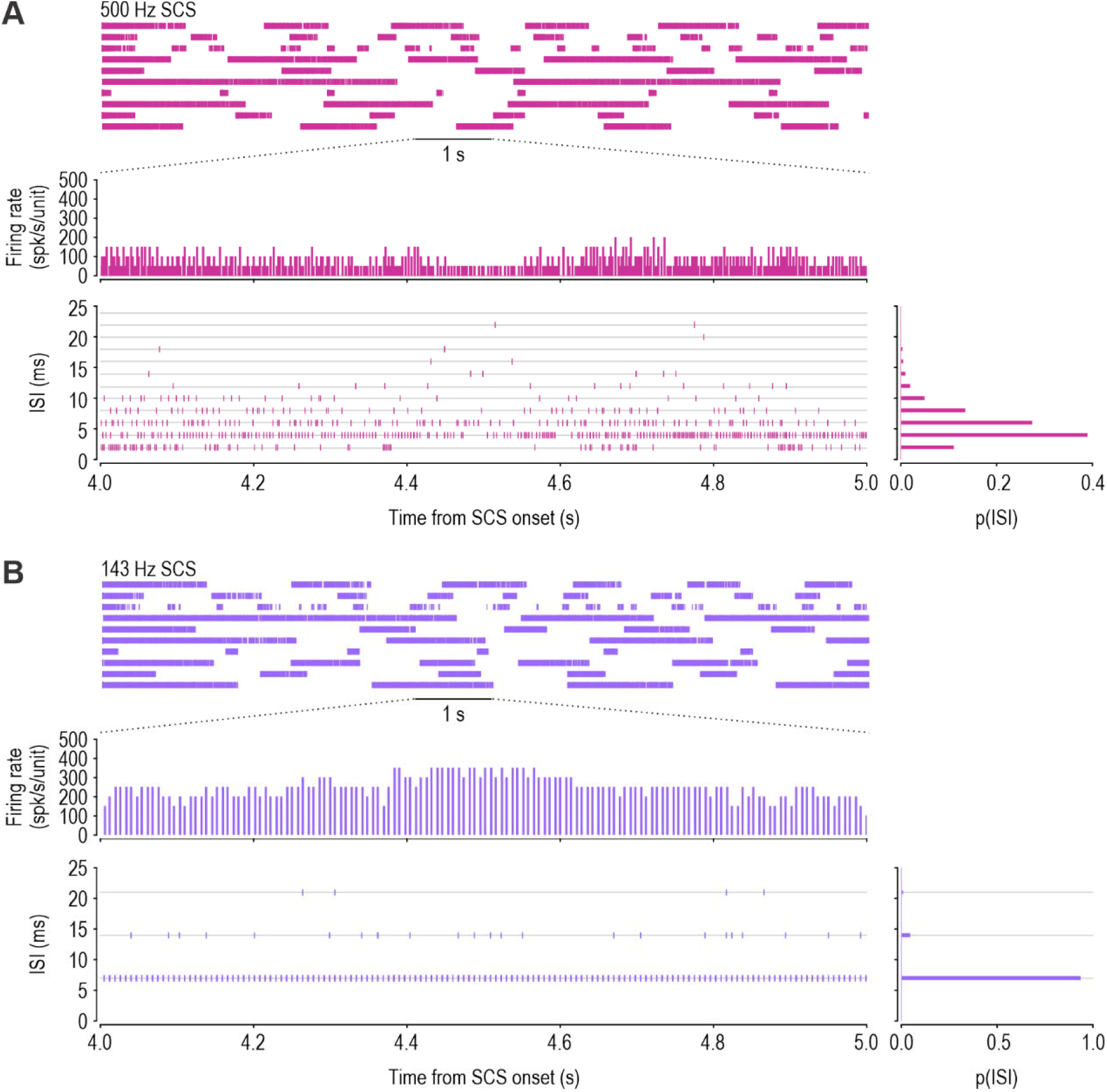
Testing of phenomenological model at additional SCS frequencies. Based on a refractory period of ∼3 ms, we predicted that overdrive synchronization should occur for SCS frequencies >333 Hz (=1/3 ms). To test this, we simulated two additional SCS frequencies. **(A)** SCS at 500 Hz evoked asynchronous spiking, consistent with our prediction and with experimental data in Supplementary Figure 7B. **(B)** SCS at 143 Hz evoked synchronous spiking, consistent with our prediction and with data in Figure 5.

**Supplementary Figure 11.**
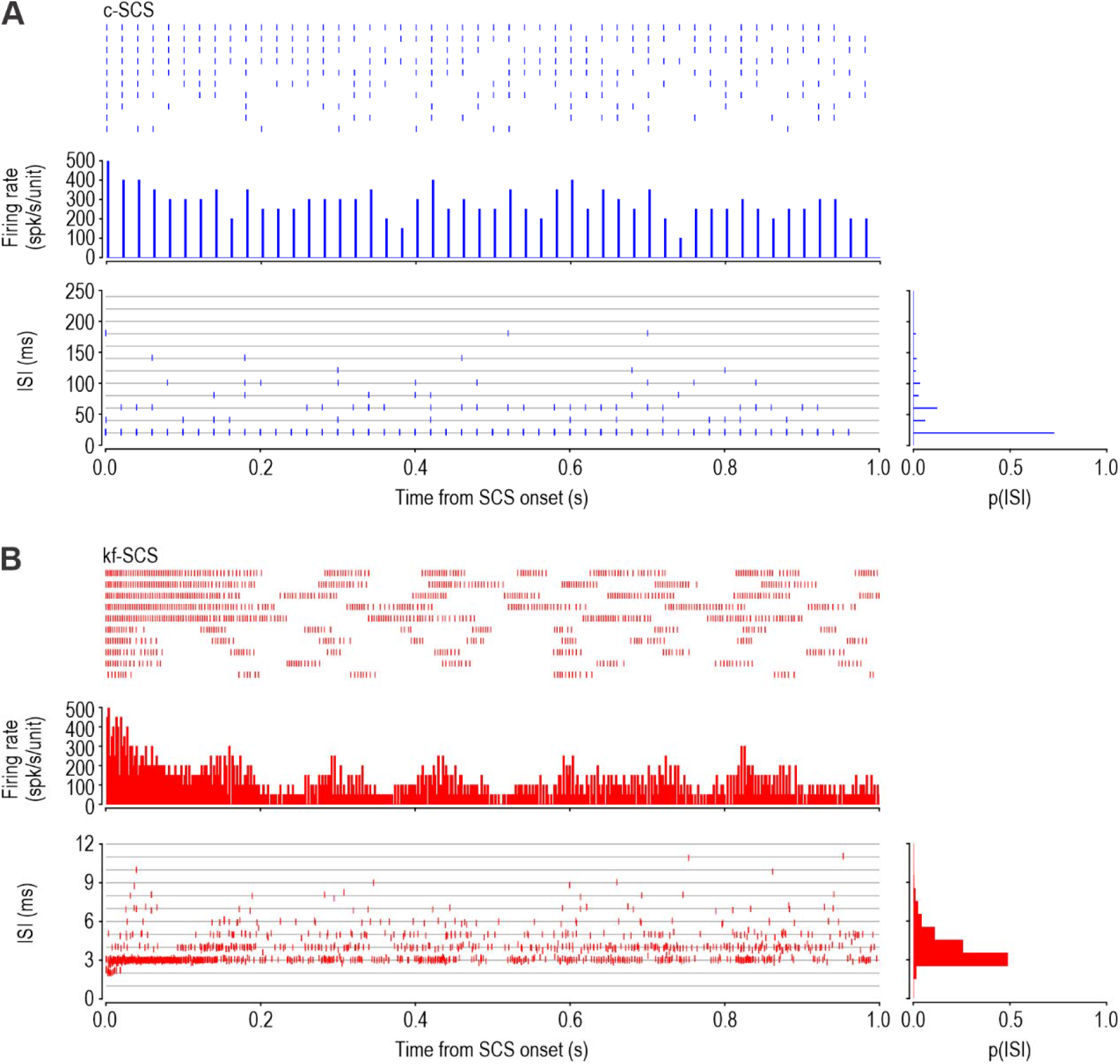
Conductance-based human axon models respond to c-SCS and kf-SCS with synchronous and asynchronous spiking, respectively. Data here are from the same simulations shown in Figure 4D, but are analyzed more fully. The same pulse amplitude and waveform were used for c-SCS and kf-SCS. Each axon model experienced different noise and had slight variations in its sodium conductance density (to simulate cellular heterogeneity) and stimulus amplitude (to simulate variable exposure to the electric field). See Methods for details. **(A)** During c-SCS, most spikes occurred at the 20 ms interpulse interval, resulting in synchronized spiking across the population. **(B)** During kf-SCS, irregular skipping caused most spikes to occur with an interval between 2 and 6 ms, resulting in asynchronous spiking.

**Supplementary Figure 12.**
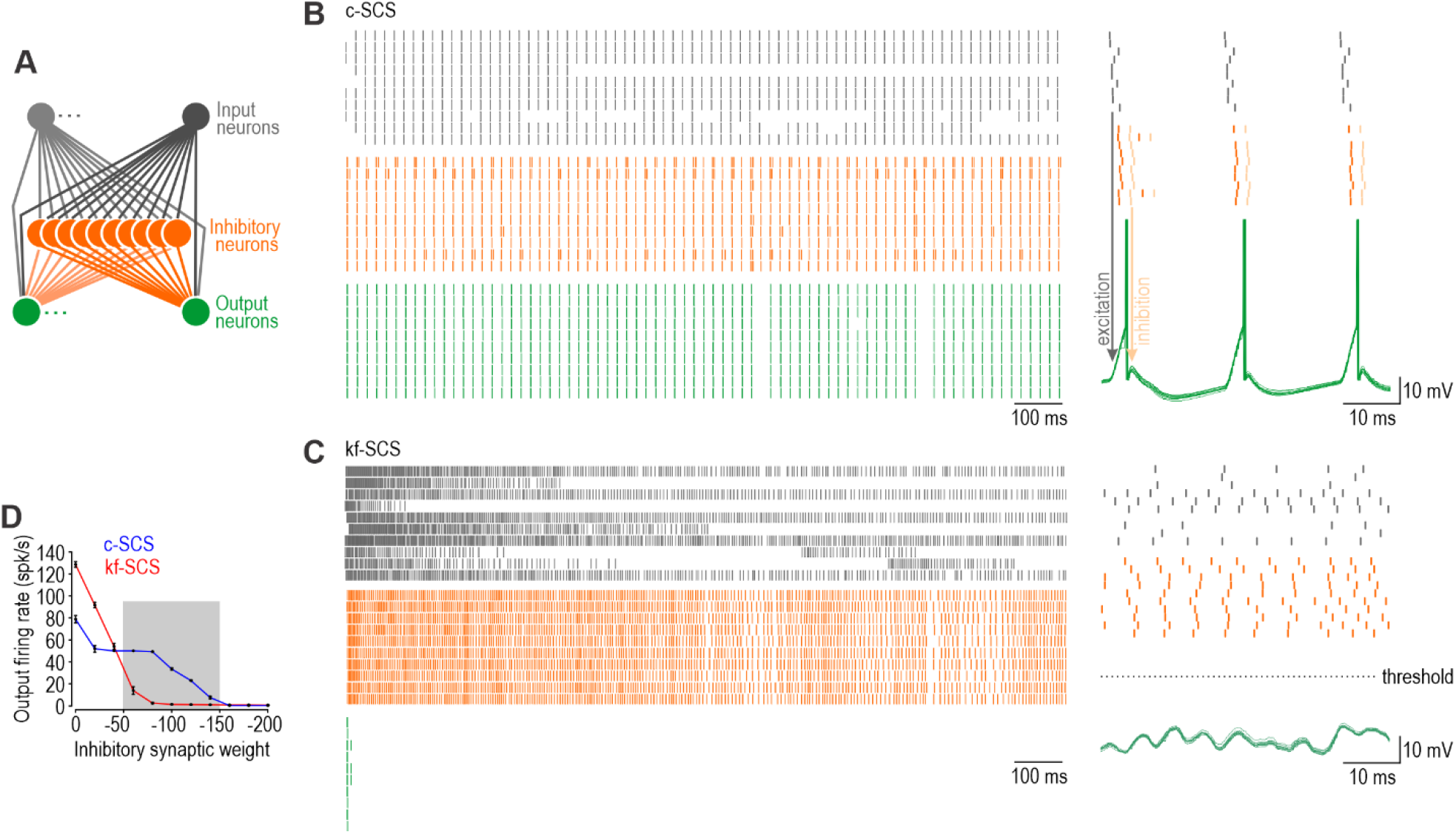
Differential blockade of synchronous and asynchronous spikes by feedforward inhibition. **(A)** Cartoon of feedforward inhibitory circuit comprising 10 input neurons (grey), 10 inhibitory neurons (orange), and 10 output neurons (green). Inhibitory and output neurons were simulated as leaky integrate-and-fire models while input neurons each provided one of 10 spike trains recorded during c-SCS or kf-SCS selected from Figure 2A to produce a median firing rate of ∼50 spk/s in both conditions. See Methods for details. **(B)** During c-SCS, each volley of synchronous excitatory inputs activates inhibitory neurons and output neurons. Output neurons reach threshold and fire before the arrival of inhibitory inputs, which arrive 2 ms after the inhibitory neuron fires (delay depicted as shift from orange to gold spikes in enlarged view on right). Under these conditions, output neurons fire synchronously at 50 spk/s. **(C)** During kf-SCS, asynchronous excitatory inputs depolarize output neurons more slowly, allowing inhibitory inputs, which are also less synchronized, to prevent output neurons from spiking. Output neurons fire only at the onset of kf-SCS. **(D)** Summary of output neuron firing rate as a function of inhibitory synaptic weight. Gray shading shows range associated with preferential blockade of asynchronous inputs; panels B and C show responses for inhibitory weight of -80. Asynchronous spikes are more readily blocked than synchronous spikes by feedforward inhibition, unless inhibition is particularly weak. Feedforward inhibition could occur at multiple points along the pathway conveying somatosensory information to the brain, including the first relay points in the spinal dorsal horn and dorsal column nuclei^74^. Absence of cortical activation during kf-SCS (see Fig 1D) suggests that kf-SCS-evoked signals are blocked subcortically, which would also explain the attenuation of SEPs and the H reflex (see Figs. 1C and B) whose triggering was temporally independent of the SCS pulses. Temporal relationships are important insofar as c-SCS-evoked input volleys evoke intermittent (as opposed to gap-free) inhibition and arrive at the weakest phase of that inhibition.

**Supplementary Figure 13.**
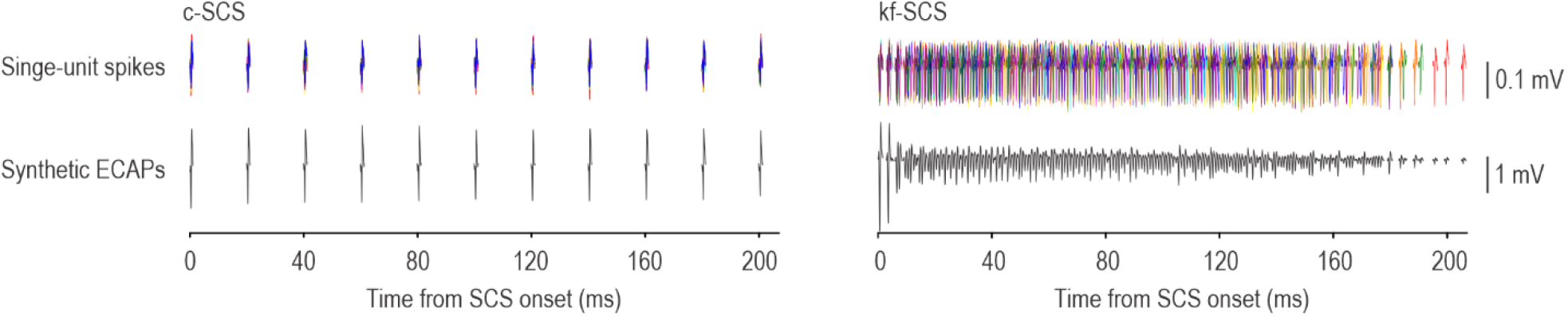
Summation of single-units spikes to model ECAPs. ECAPs were modeled here by summing the voltage waveforms for single-unit spikes extracted from 20 epochs (different colors). Each epoch represents a different burst recorded from one neuron; the first spike from each epoch was aligned to mimic synchrony at the onset of SCS. Data from different neurons (from different rats) are depicted for c-SCS and kf-SCS. During c-SCS (left), 1:1 entrainment of single-unit spikes to SCS pulses leads to large ECAPs. During kf-SCS (right), irregular skipping results in poorly synchronized spiking after the first two spikes, which in turns leads to small and irregular ECAPs. Despite the absence of cellular heterogeneity (epochs for each type of SCS are from the same neuron), uncorrelated noise was sufficient to desynchronize spiking and severely diminish ECAP amplitude. Note that single-unit spikes depicted here, based on a recording inside the DRG, are about 100× larger than for an epidural recording over the DC. In typical (clinical) ECAP recordings, many more single-units spikes must summate to produce sizeable ECAPs (see text).

